# The small G-protein Rac1 in the dorsomedial striatum promotes alcohol-dependent structural plasticity and goal-directed learning in mice

**DOI:** 10.1101/2023.08.30.555562

**Authors:** Zachary W. Hoisington, Alexandra Salvi, Sophie Laguesse, Yann Ehinger, Chhavi Shukla, Khanhky Phamluong, Dorit Ron

**Affiliations:** Alcohol and Addiction Research Group, Department of Neurology, University of California San Francisco, USA, 94107; GIGA-Stem Cells and GIGA-Neurosciences, Interdisciplinary Cluster for Applied Genoproteomics (GIGA-R), University of Liège, Belgium

## Abstract

The small G-protein Rac1 promotes the formation of filamentous actin (F-Actin). Actin is a major component of dendritic spines, and we previously found that alcohol alters actin composition and dendritic spine structure in the nucleus accumbens (NAc) and the dorsomedial striatum (DMS). To examine if Rac1 contributes to these alcohol-mediated adaptations, we measured the level of GTP-bound active Rac1 in the striatum of mice following 7 weeks of intermittent access to 20% alcohol. We found that chronic alcohol intake activates Rac1 in the DMS of male mice. In contrast, Rac1 is not activated by alcohol in the NAc and DLS of male mice, or in the DMS of female mice. Similarly, closely related small G-proteins are not activated by alcohol in the DMS, and Rac1 activity is not increased in the DMS by moderate alcohol or natural reward. To determine the consequences of alcohol-dependent Rac1 activation in the DMS of male mice, we inhibited endogenous Rac1 by infecting the DMS of mice with an AAV expressing a dominant negative form of the small G-protein (Rac1-DN). We found that overexpression of AAV-Rac1-DN in the DMS inhibits alcohol-mediated Rac1 signaling and attenuates alcohol-mediated F-actin polymerization, which corresponded with a decrease in dendritic arborization and spine maturation. Finally, we provide evidence to suggest that Rac1 in the DMS plays a role in alcohol-associated goal-directed learning. Together, our data suggest that Rac1 in the DMS plays an important role in alcohol-dependent structural plasticity and aberrant learning.

**Significance Statement:** Addiction, including alcohol use disorder, is characterized by molecular and cellular adaptations that promote maladaptive behaviors. We found that Rac1 was activated by alcohol in the dorsomedial striatum (DMS) of male mice. We show that alcohol-mediated Rac1 signaling is responsible for alterations in actin dynamics and neuronal morphology. We also present data to suggest that Rac1 is important for alcohol-associated learning processes. These results suggest that Rac1 in the DMS is an important contributor to adaptations that promote alcohol use disorder.

## Introduction

Rac1 (Ras-related C3 botulinum toxin substrate 1) is a small G-protein belonging to the Rho family of GTPases (Van Aelst and D’Souza-Schorey, 1997; Vetter and Wittinghofer, 2001; Bosco et al., 2009). Rac1 is expressed ubiquitously and plays a role in processes such as actin polymerization, endocytosis, transcription, and cell growth (Nakayama et al., 2000; Ridley, 2006; Bosco et al., 2009). Rac1 is highly expressed in the central nervous system (CNS) (Corbetta et al., 2009). Rac1 cycles between a GDP-bound inactive state and a GTP-bound active state (Vetter and Wittinghofer, 2001). In the CNS, the transition between GDP-bound to GTP-bound Rac1 is catalyzed by the guanine nucleotide-exchange factors (GEFs) Tiam1 and Karilin-7 (Vetter and Wittinghofer, 2001; Tolias et al., 2005; Xie et al., 2007). The Rac1 GTPase activating proteins (GAPs), such as RacGAP1 or FilGAP (Toure et al., 1998; Nguyen et al., 2018), initiate the intrinsic GTPase activity of Rac1 resulting in the conversion of GTP to GDP (Vetter and Wittinghofer, 2001). Upon activation, Rac1 binds to p21-activated kinase (PAK1) leading to PAK1 autophosphorylation and activation (Bokoch, 2003). PAK1 phosphorylates and activates LIM kinase (LIMK) (Edwards et al., 1999). LIMK then phosphorylates cofilin (Yang et al., 1998; Bernard, 2007; Scott and Olson, 2007). Cofilin’s function is to sever actin filaments (F-actin) into globular actin (G-actin) (Yang et al., 1998; Bamburg, 1999; Maciver and Hussey, 2002; Andrianantoandro and Pollard, 2006; Chin et al., 2016; Kanellos and Frame, 2016). LIMK phosphorylation of cofilin prevents cofilin’s ability to cleave actin, therefore enabling the formation and maintenance of F-actin (Yang et al., 1998; Pavlov et al., 2007; Scott and Olson, 2007). This mechanism promotes spine enlargement and stabilizes and strengthens synapses (Honkura et al., 2008; Chazeau and Giannone, 2016; Costa et al., 2020). As such, Rac1 is involved in long-term potentiation, the cellular mechanism of learning and memory (Haditsch et al., 2009; Haditsch et al., 2013; Lv et al., 2019).

Malfunction of Rac1 has been associated with multiple neurological and psychiatric disorders. For example, abnormal expression of Rac1 has been observed in humans with schizophrenia, autism spectrum disorders, and fragile X syndrome (Tejada-Simon, 2015; Reijnders et al., 2017; Wang et al., 2020). A reduction of Rac1 expression is also associated with stress, depression, and anhedonia in mice (Golden et al., 2013), symptoms that often coincide with addiction (Koob and Kreek, 2007). Abnormal Rac1 function has been linked to drugs of abuse. Specifically, Rac1 activity is inhibited after repeated, but not acute, cocaine administration in the nucleus accumbens (NAc) of mice (Dietz et al., 2012), and Rac1-dependent mechanisms affect the extinction of aversive opiate withdrawal memories (Wang et al., 2017). Finally, Rac1 orthologs have been shown to regulate acute alcohol sensitivity in *Drosophila melanogaster* (Peru et al., 2012). However, Rac1’s function in alcohol use disorder (AUD) has not been investigated in a mammalian system.

Previously, we found that excessive chronic alcohol consumption promotes the formation of F-actin in the NAc (Laguesse et al., 2017) and in the dorsomedial striatum (DMS) of mice (Wang et al., 2015; Laguesse et al., 2018). We further found that heavy alcohol consumption in mice results in a structural remodeling in both brain regions, leading to the maturation of dendritic spines in the NAc and in the branching of dendrites and remodeling of dendritic spines in DMS (Wang et al., 2015; Laguesse et al., 2018). Here, we examined the possibility that Rac1 controls these alcohol-dependent structural adaptations in the NAc and/or the DMS.

## Materials and Methods

### Antibodies

Rabbit anti-pLIMK (#ab38508) antibodies were purchased from Abcam (Waltham, MA). Chicken anti-GFP (#A10262) antibodies were purchased from Life Technologies (Carlsbad, CA). Rabbit anti-LIMK1 (#3842S), Cofilin (#3312S), pCofilin Ser3 (#3311S), RhoA (#6789S), Cdc42 (#2246S) antibodies were purchased from Cell signaling technology (Danvers, MA). Mouse anti-GAPDH (#G8795) antibodies, anti-actin (#A2228) antibodies, phosphatase inhibitor Cocktails 2 and 3 were from Sigma Aldrich (St. Louis, MO). Mouse anti-Rac1 antibodies (#ARC03), the G-Actin/F-Actin assay kit (BK037), the Rac1 pull-down activation assay kit (#BK035), the RhoA pull-down activation assay kit (#BK036), and the Cdc42 pull-down activation assay kit (#BK034) were purchased from Cytoskeleton, Inc. (Denver, CO). Nitrocellulose membranes and Enhance Chemiluminescence (ECL) were purchased from Millipore (Burlington, MA). EDTA-free complete mini–Protease Inhibitor Cocktails were from Roche (Indianapolis, IN). Pierce bicinchoninic acid (BCA) protein assay kit was purchased from Thermo Scientific (Rockford, IL). Ethyl alcohol (190 proof) was purchased from VWR (Radnor, PA). NuPAGE Bis-Tris precast gels were purchased from Life Technologies (Carlsbad, CA). Donkey anti-rabbit horseradish peroxidase (HRP) and donkey anti-mouse HRP conjugated secondary antibodies were purchased from Jackson ImmunoResearch (West Grove, PA). The donkey anti-mouse IgG AlexaFluor 564 and donkey anti-chicken AlexaFluor 488 antibodies were purchased from Life Technologies (Carlsbad, CA).

### Animals

Male and female C57BL/6J mice (Jackson Laboratory, Bar Harbor, ME) arrived at 6-8 weeks old and were allowed one week of habituation before experiments began. Mice were housed individually in a reversed 12-hour light-dark cycle room (lights on at 22:00/off at 10:00) that was temperature- and humidity-controlled. Unrestricted amounts of food and water were provided. Animal procedures were approved by the UCSF Institutional Animal Care and Use Committee (IACUC).

### Preparation of solutions

Alcohol solution was prepared from an absolute anhydrous alcohol solution (190 proof) diluted to 20% (v/v) in tap water for alcohol-drinking experiments. Sucrose solution (1%) was dissolved in tap water (w/v).

### Collection of brain samples for biochemical analyses

Mice were euthanized 4 hours after the beginning of the last drinking session (“binge” timepoint) or 24 hours after the last drinking session (“withdrawal” timepoint). Brains were removed and dissected on an ice-cold platform into 1mm sections, and specific subregions were dissected based on Allen Brain Atlas anatomy.

### Western blot analysis

Tissue was homogenized in ice-cold radio immunoprecipitation assay (RIPA) buffer (in mM: 50 Tris-HCL, 5 EDTA, 120 NaCl, and 1%NP-40, 0.1% deoxycholate, 0.5% SDS, proteases and phosphatases inhibitors). Samples were homogenized using a sonic dismembrator. Protein content was determined using BCA™ protein assay kit. Thirty µg of each tissue lysate was loaded for separation by SDS-PAGE then transferred onto nitrocellulose membrane at 300mA for 2 hours. Membranes were blocked with 5% milk-PBS containing 0.1% Tween 20 at room temperature (RT) for 30 minutes and then probed with the appropriated primary antibodies overnight at 4°C. The membranes were washed and probed with HRP-conjugated secondary antibodies for one hour at room temperature. Membranes were developed using the enhanced chemiluminescent reagent (ECL) and band intensities were quantified using ImageJ software (NIH).

### Small G-protein activation assay

Small G-protein activity was determined using the Rac1, RhoA, or Cdc42 activation *in vivo* assay biochemical kits for the respective protein (Cytoskeleton Inc., Denver, CO). The tissues were lysed in lysis buffer (50mM Tris-HCl pH7.4, 450mM NaCl, 1% Triton X-100) containing protease and phosphatase inhibitor cocktail. Thirty µg of each lysate was incubated with 10 µL PAK Rac1/Cdc42 binding domain (PAK-PBD)-agarose beads for Rac1 and Cdc42, or 15 µL Rhotekin Rho binding domain (Rhotekin-RBD)-agarose beads for RhoA, for 16 hours. For the control, the same amount of lysate was incubated with GDP or GTP for 15 minutes followed by incubation with PAK-PBD or Rhotekin-RBD-agarose beads for 16 hours. The beads were washed twice with washing buffer followed by boiling in 30 µL 2X sample loading buffer. The samples were analyzed by western blot.

### F-actin/G-actin assay

F-actin and G-actin content was measured using the G-actin/F-actin in vivo assay biochemical kit (Cytoskeleton Inc., Denver, CO) as previously described in (Laguesse et al., 2018) with slight modification. Tissue punches were homogenized in 250 µL cold LAS02 buffer containing protease and phosphatases inhibitors and centrifuged at 350g for 5 minutes at 4°C to remove cellular debris. Protein concentrations were determined in the supernatant using BCA™ protein assay kit. Supernatant was centrifuged at 15,000g for 30 minutes at 4°C and the new supernatant contained soluble globular actin (G-actin). The insoluble filamentous actin (F-actin) in the pellet was then resuspended and incubated on ice for 1 hour in 250 µL F-actin depolymerization buffer, with gently mixing every 15 minutes. Samples were centrifuged at 15,000g for 30 minutes at 4°C and this supernatant was used to measure F-actin content by western blot. Twenty µL of the G-actin fraction and 40 µL of the F-actin fractions were loaded and analyzed by western blot.

### Adeno-associated viruses

AAV2-Rac1-DN-GFP (AAV-Rac1-DN; 3.5×10^12^ vg/ml) was produced by the Duke Viral Vector Core (Durham, North Carolina). The plasmid containing the T17N Rac1 mutation (plasmid #22784, pCyPet-Rac1(T17N)) and AAV-GFP (pAAV.CMV.PI.EGFP.WPRE.bGH; 1.6×10^13^ vg/ml) were purchased from Addgene. AAV-GFP was diluted to match the concentration of AAV-Rac1-DN.

### Confirmation of AAV-Rac1-DN in cells

HEK293 cells were plated at 120,000 cells per well on a 12-well-plate. The media was changed to 1% FBS-DMEM 24 hours prior to the infection. The cells were then infected with 2 µL of AAV-Rac1-DN virus (3.5×10^12^ vg/ml). Seventy-two hours after the infection, cells were incubated with 10% FBS for 30 minutes. The cells were lysed and analyzed by western blot.

### Stereotaxic surgery

Mice were anesthetized using vaporized isoflurane and were fixed in a stereotaxic frame (David Kopf Instruments). To overexpress the virus in the entirety of the DMS, bilateral microinfusions were accomplished using stainless steel injectors (33 gauge; Small Parts Incorporated) connected to Hamilton syringes (10 µL, 1701) at two sites in the DMS (anteroposterior (AP) +1.1 mm, mediolateral (ML) ± 1.2 mm, dorsoventral (DV) -3 mm and AP +0.66 mm, ML ± 1.2 mm, DV -2.8 mm measured from bregma). Animals received a 1μL bilateral infusion of AAV-GFP or AAV-Rac1-DN-GFP (3.5×10^12^ vg/ml) at an infusion rate of 0.1μL/min controlled by an automatic pump (Harvard Apparatus). The injectors were left at the infusion site for 10 minutes after the conclusion of the infusion to allow the virus to diffuse.

To image single neurons, a low-titer (7×10^9^ - 3.5×10^10^ vg/ml) of AAV-GFP or AAV-Rac1-DN-GFP was infused into the DMS (AP +1.1 mm, ML ± 1.2 mm, DV -2.8 mm). 0.5μL of each virus was infused bilaterally at a rate of 0.1μL/min. Injectors were left in place for 10 minutes for viral diffusion.

### *In vivo* confirmation of viral expression

After the conclusion of an experiment, animals were euthanized by cervical dislocation and the brains were removed. The brain was dissected on ice into 1-mm-thick coronal sections and green fluorescent protein (GFP) was visualized and imaged using an EVOS FL tabletop fluorescent microscope (ThermoFisher Scientific, Waltham, MA). Mice with surgeries that failed to elicit viral overexpression were excluded from the study.

### Immunostaining

Mice were euthanized by transcardial perfusion with 0.01M PBS followed by 4% paraformaldehyde (PFA) in phosphate buffer, for 5 minutes each step. Brains were removed and postfixed in 4% PFA for 24 hours before being transferred to a 30% sucrose solution for 3 days. Brains were then frozen rapidly on dry ice before being coronally sectioned into 100 μm sections using a Leica CM3050 cryostat (Leica Biosystems, Richmond, IL). Slices were placed in 0.01M PBS and the ones containing the site of infection were selected to be stained. Sections were blocked with 5% normal donkey serum for 4 hours. Slices were incubated in the primary antibody over 48 hours at 4°C on an orbital shaker. After primary antibody incubation, slices were washed 3 times for 45 minutes each in PBS before secondary antibody incubation overnight at 4°C on an orbital shaker. There was another cycle of washing before placing the slices onto Superfrost Plus microscope slides (Fisher Scientific, Hampton, NH) and mounting slides using Prolong Gold mounting medium (ThermoFisher Scientific, Waltham, MA). Slides were allowed 24 hours to dry before the edges were sealed to prevent dehydration.

### Morphological analysis

Low titer of AAV-GFP or AAV-Rac1-DN (7×10^9^ - 3.5×10^10^ vg/ml) was infused bilaterally into the DMS. After 1 week of recovery, mice underwent 7 weeks of IA20%2BC. Twenty-four hours after the last drinking session, mice were euthanized, perfused, processed, and 100 µm coronal sections were collected. Images of GFP-stained DMS neurons were acquired with a spinning disk confocal microscope with a 40x objective and a Z-interval of 1μm (Nikon, Tokyo, Japan). Images were deconvoluted using AutoQuantX (v3.1.3, Media cybernetics, Inc, Rockville, MD), and GFP neurons were traced using Neurolucida software (MBF Biosciences, Williston, VT). *Scholl analysis*: Dendritic branches were quantified using Sholl analysis (Sholl, 1953). Starting radius of dendritic branches was 10μm and end radius was 160μm from the center of the soma with an interval of 10μm between radii. Cell body area was measured using Fiji software (NIH). *Spine Analysis:* Images of GFP-stained DMS neurons were acquired with a spinning disk confocal microscope with a 100x objective and a Z-interval of 0.1μm (Nikon, Tokyo, Japan). Only distal dendrites (3^rd^ or 4^th^ order) were analyzed. Morphological properties were analyzed using Imaris software (Oxford Instruments, Concord, MA). Dendritic spines were classified based on a combination of length and head and neck morphology. Spines were classified into four subtypes as in (Laguesse et al., 2018): filipodia (>2μm length, indiscernible head), stubby (<1μm length, no visible neck), mushroom (>0.4μm head width, short neck), thin (head width <0.4μm, long neck).

### Behavioral paradigms

2-bottle choice drinking paradigms

#### Intermittent access to 20% alcohol (IA20%2BC)

The paradigm was conducted as described in (Laguesse et al., 2017). Mice were given 24-hour access to one bottle of 20% alcohol (v/v) in tap water or one bottle of water. Drinking sessions started at 11:30AM Monday, Wednesday, and Friday with 24- or 48-hour (weekend) withdrawal periods in between during which only water was available. The bottles were alternated each session to prevent a side preference from developing. The bottles were weighed at the end of each session. Bottles of 20% alcohol and water were also placed on an empty cage. The change in weight of these bottles during the session was used to quantify spillage. This weight alteration was deducted from the weight change observed in the bottles assigned to the animals to adjust for experimental spillage. Mice were weighed once a week. Animals that drank over 7g/kg/24hr were included in the study. See **Table 1** for average values in each of the experiments. Percentile of alcohol preference (preference index) was calculated by dividing the amount of alcohol consumed to the amount of water+alcohol ×100.

**Table 1.** Average Alcohol Consumption for biochemical experiments.

| **Figure** | **Average Consumption (g/kg/24hr)** | **SEM (±g/kg/24hr)** |
| --- | --- | --- |
| 1B NAc | 13.38 | 1.025 |
| 1C DLS | 13.38 | 1.060 |
| 1D DMS | 14.53 | 0.982 |
| 1E Female DMS | 19.59 | 0.979 |
| 1F RhoA | 12.31 | 0.4461 |
| 1G Cdc42 | 13.67 | 0.7354 |
| 1H 10%CA | 8.787 | 0.2611 |
| 1J 4wk DMS | 12.560 | 1.678 |
| 1K pLIMK pCofilin | 15.34 | 0.5169 |
| 2H-M AAV-GFP | 14.00 | 0.7264 |
| 2H-M AAV-Rac1-DN | 13.68 | 0.4589 |
| 3,4 AAV-GFP | 10.63 | 0.5058 |
| 3,4 AAV-Rac1-DN | 10.88 | 1.223 |
| 6 AAV-GFP | 12.10 | 0.7463 |
| 6 AAV-Rac1-DN | 12.36 | 1.046 |
| 1I Sucrose | 174.81 (ml/kg) | 17.87 (ml/kg) |

#### Intermittent access to 1% sucrose

Mice had access to a 2-bottle choice between water and 1% sucrose three days a week for 24-hour periods for two weeks (Laguesse et al., 2017). Bottles were weighed at the end of each session and switched in between. Between sessions, only water was available. The bottle weights were spill-adjusted after each session. Mice were weighed weekly, and 1% sucrose consumption was measured in milliliters per kilogram of bodyweight (ml/kg). Food was available *ad libitum*.

#### Continuous access to 10% alcohol

Mice had access to a bottle of 10% alcohol (v/v) in tap water and a bottle of water for 24 hours a day for 3 weeks (21 drinking sessions) (Laguesse et al., 2017) Bottles were removed and weighed daily. Bottle positions were alternated to reduce side preference development. The bottle weights were spill-adjusted after each session. Mice were weighed weekly and alcohol consumption was calculated for each in grams per kilogram.

### Operant self-administration (OSA) paradigms

#### Alcohol operant self-administration

OSA was conducted as described in (Laguesse et al., 2017). Mice underwent stereotaxic surgery and IA20%2BC for 7-8 weeks as described above. Prior to training, animals were handled for 1-2 minutes per day for 3 consecutive days. OSA was conducted during the dark phase of the reverse light/dark cycle in operant chambers (length: 22 cm, width: 20 cm, height: 14 cm). OSA boxes were connected to a computer running MED-PC to control and record session activity. The operant chambers (Med-Associates; Georgia, VT) had two levers (length: 1.5 cm, distance between: 11 cm, height above floor: 2.5 cm) on one wall. The operant chambers were also equipped with a reward port between the levers (height above floor: 0.5 cm) that included a photo-beam to track port entries. Upon reward delivery, a 3-second tone (2900Hz) and a cue-light above the reward port was activated. To receive a reward, the mouse must press the active lever in a fixed-ratio 1 (FR1) schedule, where one active-lever press leads to one reward. A successful completion of this condition resulted in a delivery of 10 µL of 20% alcohol to the reward port. The mouse must enter the reward port twice to reactivate the active lever, ensuring consumption of the reward. Animals received a total of 20 hours of training time in the paradigm in the first week (two 6-hour and two 4-hour sessions) before transitioning to 2-hour afternoon sessions that began consistently at 13:00. After 8 2-hour FR1 sessions, the complexity of the task was increased to FR2, where two active lever presses were required for one reward. Active lever presses, inactive lever presses, port entries, and reward deliveries were measured. Consumption was normalized based on mouse body weight and rewards administered. Discrimination index was calculated as the percentage of active lever presses divided by total presses. The proportion of reward lever presses statistic was calculated by comparing lever presses that led to a reward to total lever presses, including inactive lever presses. Mice with low viral expression were excluded from the study.

#### Contingency degradation

The contingency degradation experiment was conducted as previously described in (Morisot et al., 2019a). Mice were first trained in the OSA boxes on an FR1 schedule of reinforcement with 20% alcohol. Mice completed three 6-hour sessions and three 4-hour sessions before transitioning to 2-hour sessions starting at 13:00. After two 2-hour sessions under the FR1 schedule, mice progressed to random ratio (RR) training. During RR OSA, rewards were delivered following a random number of lever presses. Mice completed five sessions under RR2 (one reward delivery following an average of two lever presses with number of presses ranging from one to three), followed by five sessions of RR3 (number of presses ranged from two to four), and ten sessions of RR4 (number of presses ranged three to five). After completion of training, mice underwent two types of contingency degradation testing sessions, non-degraded (ND) and degraded (D). During ND sessions, active lever pressing led to the same cue and reward delivery as RR4 training. However, during D sessions, active lever pressing had no outcome, and rewards were delivered regularly throughout the session, determined by the average reward delivery rate of the last five RR4 training sessions. One mouse was excluded due to low viral expression.

#### Sucrose operant self-administration

OSA of 1% sucrose was slightly modified from conditions described in (Laguesse et al., 2017). Specifically, mice underwent stereotaxic surgery to overexpress AAV-GFP or AAV-Rac1-DN-GFP in the DMS. After allowing for viral overexpression, mice began 1% sucrose self-administration training in the chambers previously described with two 6-hour and two 4-hour training sessions in the first week. Animals were handled for 1-2 minutes daily for 3 consecutive days before training began. They then transitioned to 2-hour afternoon sessions that began consistently at 13:00. After 8 2-hour FR1 sessions, the mice transitioned to FR2. Mice with low viral expression were excluded from the study.

## Open-field test

Mice infected with either AAV-GFP or AAV-Rac1-DN were placed in an open field (43cm x 43cm) in low-light conditions and allowed to explore for 20 minutes (Warnault et al., 2016). Locomotor activity was tracked using EthoVision XT software (Noldus, Leesburg, VA), and total movement (cm) was reported.

## Data analysis

GraphPad Prism 9 and MATLAB were used for statistical analysis.

### Biochemical analysis

Data were analyzed using the appropriate statistical test, including one-way ANOVA, two-way ANOVA, three-way ANOVA, or two-tailed t-test for normal populations, or Kruskal-Wallis for non-normal populations. *Post hoc* testing was done using Šidák’s multiple comparisons test. For data represented in Figures 3 and 4, statistical analysis was separated by independent variable.

IA20%2BC numbers are expressed as the mean ± SEM consumption over the final two weeks (**Table 1**). Data was first tested for normality using the Shapiro-Wilk normality test with accompanying QQ plot. Parametric tests were performed on data samples deemed to be derived from normal populations. The results were determined to be statistically significant if the resulting p-value was less than 0.05.

### Behavioral analysis

IA-2BC and OSA data were analyzed using a two-way repeated-measures ANOVA, followed by *post hoc* Šidák’s multiple comparisons test. A two-tailed t-test was performed on the open-field locomotion. The results were determined to be statistically significant if the resulting p-value was less than 0.05.

## Results

### Alcohol activates Rac1 in the DMS of male mice

We first tested if repeated cycles of alcohol binge drinking and withdrawal activates Rac1 in the striatum of male mice. Specifically, animals were subjected to 7 weeks of 20% alcohol or water choice for 24 hours a day, 3 days a week, with 24- or 48- (weekend) hour withdrawal periods in between during which mice had access to water only. Mice consuming water for the same duration were used as controls (**Figure 1A**). Striatal regions were dissected and harvested 4 hours after the beginning of a drinking session (“binge”) or 24 hours after the end of a drinking session (“withdrawal”) (**Figure 1A**). Rac1-GTP pulldown assay was utilized to analyze the level of active GTP-bound vs. inactive GDP-bound Rac1. We found that the activity of Rac1 was unaltered by alcohol in the NAc (**Figure 1B**, **Table 1**) and the DLS (**Figure 1C**, **Table 1**) (NAc Kruskal-Wallis test *H* = 0.5731, p = 0.7761; DLS One-way ANOVA: F_(2,15)_ = 0.0139, p = 0.9862). In contrast, we discovered that 7 weeks of IA20%2BC produced a robust activation of Rac1 in the DMS during both binge and withdrawal (**Figure 1D**, **Table 1**) (One-way ANOVA: F_(2, 15)_ = 8.233, p = 0.0039; *Post hoc*: Water vs. Binge p = 0.0058, W vs. WD p = 0.0068). The same pattern of Rac1 activation was also observed during withdrawal after 4 weeks of IA20%2BC (**Figure 1J**, **Table 1**) (Unpaired t-test: *t*_(6)_ = 2,938, p = 0.0260). In contrast, Rac1 activity was unaltered in the DMS of female mice undergoing 7 weeks of IA20%2BC (**Figure 1E**, **Table 1**) (Kruskal-Wallis test *H* = 0.2456, p = 0.8968), suggesting that there are sex differences in alcohol-dependent Rac1 activation. Together, these data suggest that chronic voluntary drinking of alcohol produces a long-lasting activation of Rac1 in the DMS of male mice.

**Figure 1.**
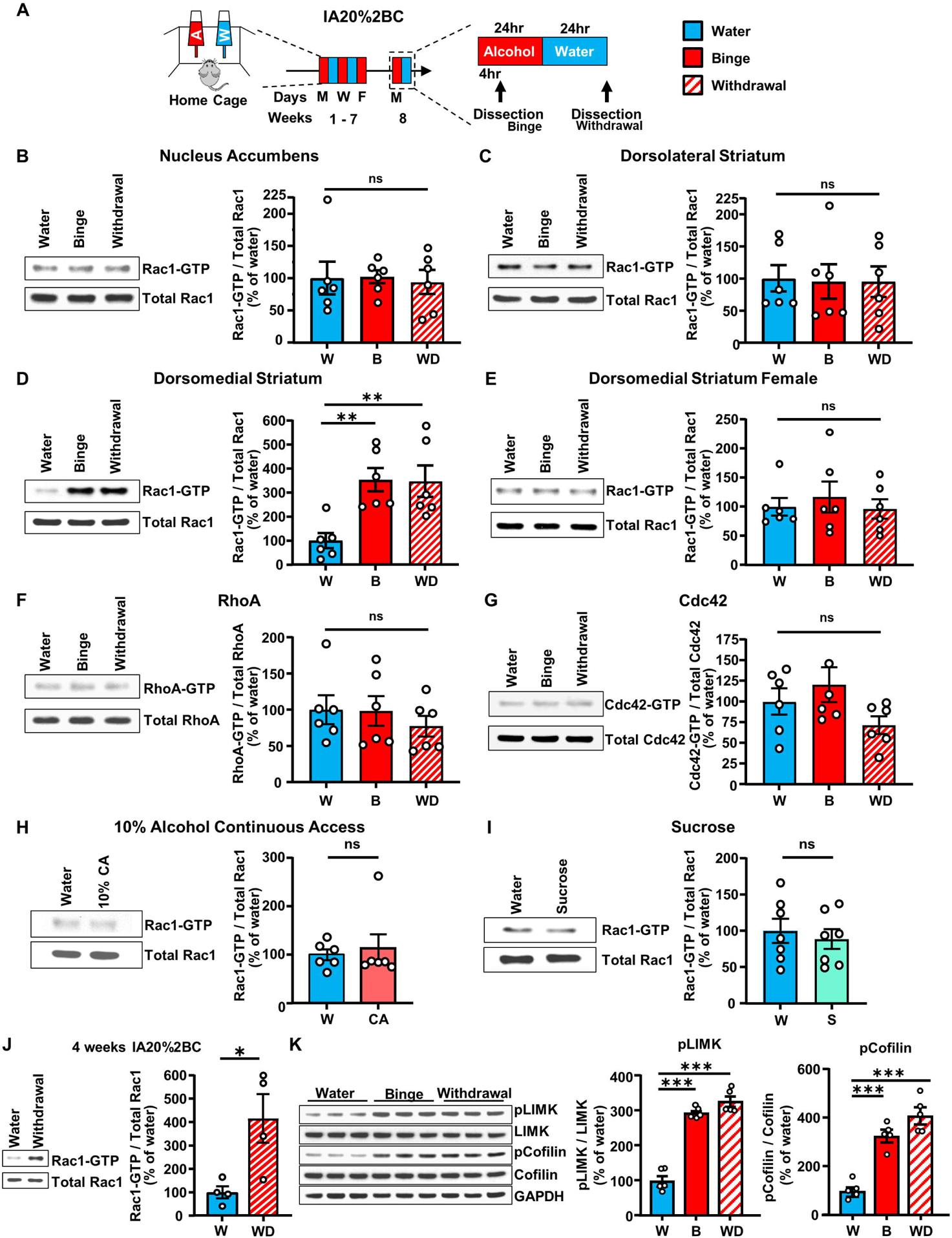
Alcohol activates Rac1 specifically in the DMS of male mice. **(A)** Intermittent access to 20% alcohol in 2-bottle-choice (IA20%2BC) experimental timeline. **(B-D)** The NAc **(B)**, the DLS **(C)**, and the DMS **(D)** were harvested 4 hours into last drinking session (binge, B) or after 24 hours of abstinence (withdrawal, WD). Rac1-GTP pull-down assay was conducted on cell lysates. Levels of GTP-bound Rac1 were normalized to total Rac1 and presented as percentage of the average of the water values. **(E)** Female mice underwent IA20%2BC before the DMS was harvested and the percentage of Rac1-GTP was calculated. RhoA-GTP **(F)** or Cdc42-GTP **(G)** pull-down assay was conducted on cell lysates after IA20%2BC and quantified using western blot. Levels of GTP-bound RhoA or Cdc42 were normalized to total respective protein and presented as percentage of average of the water values. **(H)** Mice underwent 21 sessions of 10%CA and the DMS was harvested. Rac1-GTP pull-down assay was conducted on cell lysates and quantified using western blot. Rac1-GTP was normalized to total Rac1 and presented as percentage of average of water values. **(I)** Mice underwent intermittent access to 1% sucrose for 2 weeks and the DMS was harvested. Rac1-GTP pull-down assay was performed on cell lysates, and Rac1-GTP was normalized to total Rac1 and presented as average of water values. **(J)** After 4 weeks of IA20%2BC, the DMS was harvested and Rac1-GTP pull-down assay was conducted on cell lysates. Levels of GTP-bound Rac1 were normalized to total Rac1 and presented as percentage of the average of the water values. Data are represented as mean ± SEM and analyzed with an unpaired two-tailed t-test. *p < 0.05. n = 4 per group. **(K)** Mice underwent 7 weeks of IA20%2BC and LIMK and cofilin phosphorylation in the DMS were examined using western blot analysis. The levels of phospho-LIMK (pLIMK) and phospho-Cofilin (pCofilin) were normalized to total respective protein and quantified as a percentage of the average of the water values. Data are represented as mean ± SEM and analyzed using one-way ANOVA (C,D,F, K) with *post hoc* Šidák’s multiple comparisons test, Kruskal-Wallis test (B, E, G), or unpaired two-tailed t-test (H,I,J). *p < 0.05, **p<0.01, ***p < 0.001; ns, non-significant. n = 5-7 per group.

### Rac1 activation by alcohol in the DMS is specific

Next, we set to determine the specificity of alcohol-dependent activation of Rac1 in the DMS. The closely related small G-proteins RhoA and Cdc42 have also been linked with synaptic and structural plasticity (Francis et al., 2019; Zhang et al., 2021). We found that levels of RhoA and Cdc42 bound to GTP (**Figure 1F**, **Figure 1G**, **Table 1**) were unchanged in the DMS at both binge and withdrawal timepoints as compared to water-only drinking mice (RhoA One-way ANOVA: F_(2, 15)_ = 0.472, p = 0.6323; Cdc42 Kruskal-Wallis test *H* = 3.310, p = 0.1962). To measure if Rac1 is activated in response to moderate consumption of alcohol, we exposed mice to a 10% continuous access (10%CA) 2BC alcohol drinking paradigm in which mice were allowed to choose between 10% alcohol and water continuously for 21 days, matching the number IA20%2BC sessions. We detected no change in the activation of Rac1 in the DMS after 10%CA (**Figure 1H**, **Table 1**) (Unpaired t-test: *t*_(10)_ = 0.3960, p = 0.7004), which implies that higher concentration of alcohol and/or repeated cycles of binge and withdrawal are necessary for the alcohol-dependent activation of Rac1 in the DMS.

To examine if Rac1 is activated in the DMS by a naturally rewarding substance, mice underwent intermittent access to a 1% sucrose 2BC paradigm for 2 weeks. We found that Rac1 is not activated in the DMS of sucrose drinking mice (**Figure 1I**, **Table 1**) (Unpaired t-test: *t*_(12)_ = 0.5345, p = 0.6028), suggesting that activation of Rac1 signaling in the DMS is specific for alcohol and is not shared with natural reward. Together, these data suggest that the activation of Rac1 in the DMS observed after chronic alcohol consumption is not generalized to other closely related small G-proteins in the Rho family, is specific to repeated cycles of binge and withdrawal of 20% alcohol and is not shared with natural reward.

### Alcohol promotes LIM kinase activation and cofilin phosphorylation

Rac1 activation leads to the downstream phosphorylation of LIMK which in turn phosphorylates cofilin (Edwards et al., 1999). Therefore, we examined whether alcohol-mediated Rac1 stimulation promotes the activation of LIMK and cofilin phosphorylation in the DMS. To test this question, we measured the level of LIMK phosphorylation, and thus activation, as well as the phosphorylation of cofilin in the DMS after 7 weeks of IA20%2BC. We found that the phosphorylation of both LIMK and cofilin were significantly increased after alcohol binge and withdrawal in comparison to animals that drank water only (**Figure 1K**, **Table 1**) (One-way ANOVA: LIMK F_(2, 12)_ = 96.40, p < 0.0001; *Post hoc*: Water vs. Binge p < 0.0001, Water vs. Withdrawal p < 0.0001; Cofilin F_(2, 12)_ = 44.14, p < 0.0001; *Post hoc*: Water vs. Binge p < 0.0001, Water vs. Withdrawal p < 0.0001). Together, these data suggest that long-term alcohol consumption activates Rac1-dependent signaling.

### Alcohol activation of the LIMK/cofilin signaling pathway in the DMS depends on Rac1

The LIMK/cofilin signaling pathway can also be activated by other small G-proteins such as RhoA/B, Cdc42, and Rac3 (Edwards et al., 1999; Mira et al., 2000; Swanson et al., 2017). Thus, to confirm that the upregulation of LIMK and cofilin phosphorylation after excessive alcohol use is indeed due to the increase in Rac1 activity, we used a dominant negative Rac1 mutant to inhibit the activity of the endogenous protein. Specifically, the dominant negative mutant of Rac1 (Rac1-DN) contains a threonine to asparagine substitution at residue 17 (Worthylake et al., 2000). The Rac1 mutant forms a tight complex with Rac1-specific GEFs, but does not allow the exchange of GDP to GTP, keeping the G-protein constantly inactive (Ridley et al., 1992; Wong et al., 2006; Zhang et al., 2016) (**Figure 2A**). Rac1-DN was packaged into an adeno-associated virus (AAV). First, to confirm the inhibitory action of Rac1-DN, HEK293 cells were infected with AAV-Rac1-DN in media containing 1% serum (**Figure 2B**). Next, cells were incubated for 30 minutes with media containing 10% FBS. As shown in **Figure 2B**, incubation of cells with 10% FBS increased LIMK phosphorylation in non-infected cells, which was not observed in cells infected with AAV-Rac1-DN. In contrast, ERK1/2 whose phosphorylation does not depend on Rac1 was increased by 10% serum in both uninfected cells and AAV-Rac1-DN infected cells. These data suggest that AAV-Rac1-DN selectively inhibits Rac1 signaling in cultured cells.

**Figure 2.**
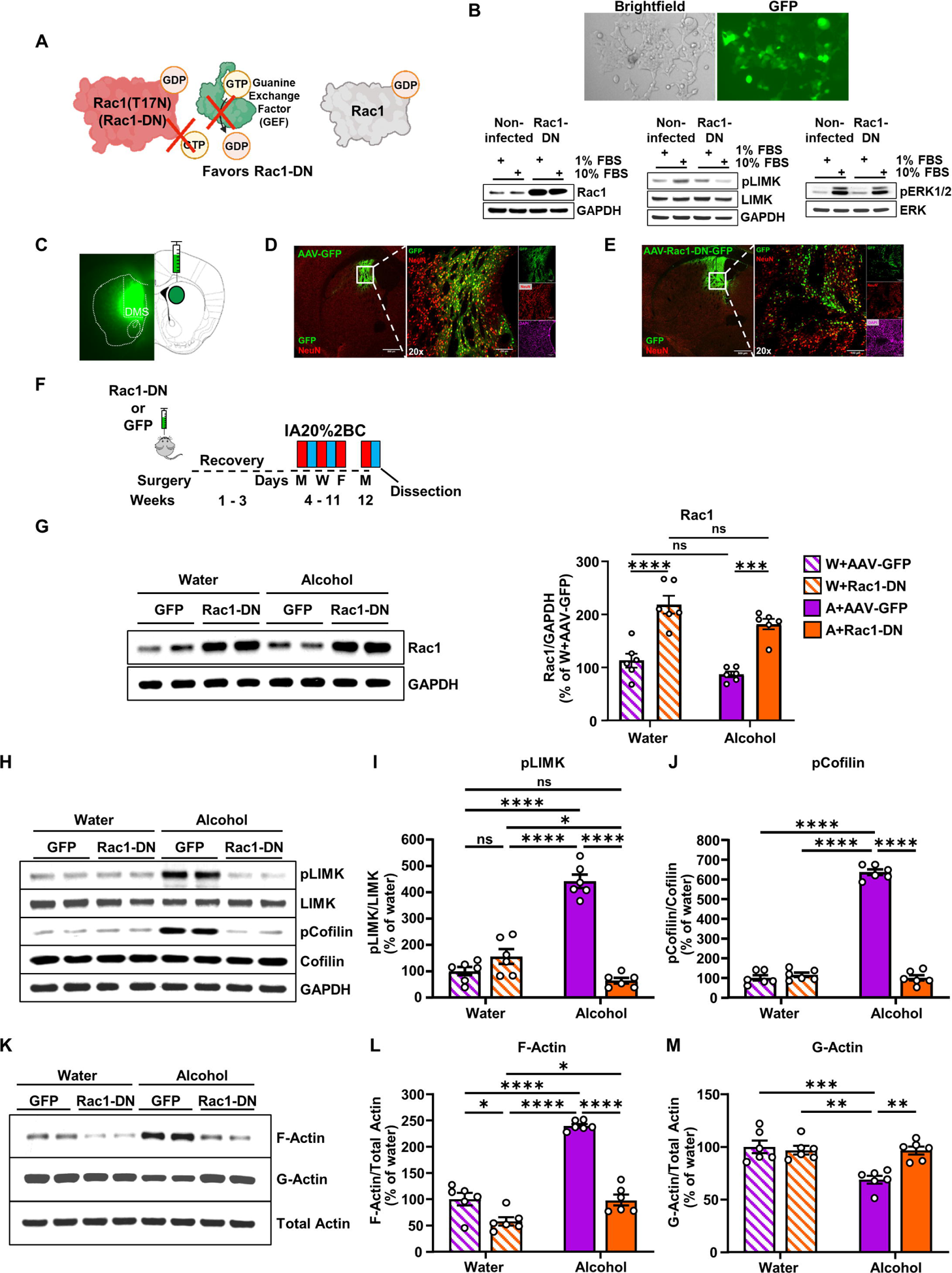
Alcohol activation of the LIMK/cofilin signaling pathway and subsequent F-actin formation in the DMS depends on Rac1. **(A)** Rac1-DN mechanism of action. Rac1-DN has a mutation in its P-loop and does not bind GTP. GEFs bind to Rac1-DN but are unable to exchange GDP for GTP. The GEFs remain bound to Rac1-DN and ignore endogenous Rac1. **(B)** HEK293 cells infected with AAV-Rac1-DN imaged in both brightfield and GFP to detect viral infection (2X magnification). Western blot analysis of Rac1 protein, phospho-LIMK, and phosphorylation of ERK1/2 in non-infected cells and cells infected with AAV-Rac1-DN after stimulation with 10% FBS. **(C)** Confirmation of viral overexpression during dissection on the DMS of a mouse infected with AAV-Rac1-DN. 2X image from EVOS FL tabletop fluorescent microscope. **(D-E)** Left images (4X magnification) depict the specificity of the infection site for AAV-GFP **(D)** and AAV-Rac1-DN **(E)**, scale bar 500μm. Right images (20X magnification) depict solely neurons infected in the DMS by both AAV-GFP **(D)** and AAV-Rac1-DN **(E)**, scale bar 100μm. Each slice is stained with anti-GFP (green) and anti-NeuN (red) antibodies, along with DAPI (magenta). **(F)** Experimental timeline. Mice received bilateral infusion of AAV-Rac1-DN or AAV-GFP in the DMS and were allowed 3 weeks for overexpression. After IA20%2BC or water only for 7 weeks the DMS was harvested. **(G)** Mice were infected with AAV-GFP or AAV-Rac1-DN before undergoing IA20%2BC. The DMS was harvested, and total Rac1 protein level was examined using western blot and normalized to GAPDH. Data are presented as mean ± SEM and analyzed using two-way ANOVA with *post hoc* Šidák’s multiple comparisons test. ***p<0.001 ****p<0.0001; ns, non-significant. n = 6 per group. **(H-J)** Phosphorylation of LIMK and cofilin were examined using western blot analysis. Levels of pLIMK and pCofilin were normalized to total respective protein and quantified as a percentage of AAV-GFP-infected, water-only animals. Data are represented as mean ± SEM and analyzed by two-way ANOVA with *post hoc* Šidák’s multiple comparisons test. *p<0.05, ****p<0.0001; ns, non-significant. n = 6 per group. **(K-M)** G-actin/F-actin assay was conducted on the DMS of mice after overexpression surgery and IA20%2BC or water-only drinking. The filamentous (F) or globular (G) actin contents were examined using western blot analysis and normalized to total actin and quantified as a percentage of AAV-GFP-infected, water-only animals. Data represented as mean ± SEM and analyzed using two-way ANOVA with Šidák’s multiple comparison. *p < 0.05, **p<0.01, ***p < 0.001, ****p<0.0001; ns, non-significant. n = 6 per group.

Next, mice were bilaterally infected with AAV-Rac1-DN (3.5×10^12^ vg/ml) or an AAV-GFP (3.5×10^12^ vg/ml) control in the DMS. As shown in **Figure 2C-E**, intense and localized viral expression was detected in mice infected with AAV-Rac1-DN. We then tested if Rac1 is required for alcohol-mediated LIMK and cofilin phosphorylation in the DMS. Three weeks after surgery, mice underwent 7 weeks of IA20%2BC before dissection (**Figure 2F**). The DMS of mice infected with AAV-GFP or AAV-Rac1-DN were exposed to the same concentration of alcohol (**Table 1**). Global Rac1 protein level was significantly increased in the DMS of mice infected with AAV-Rac1-DN compared to mice infected with AAV-GFP (**Figure 2G**) (Two-way ANOVA: Effect of virus F_(1,20)_ = 69.58, p<0.0001; Effect of alcohol F_(1,20)_ = 6.774, p = 0.0170; Effect of virus x alcohol F_(1,20)_ = 0.2164, p = 0.6468; *Post hoc* W+AAV-GFP vs. W+AAV-Rac1-DN p<0.0001; A+AAV-GFP vs. A+AAV-Rac1-DN p = 0.0001). The phosphorylation of LIMK and cofilin were measured in the DMS of water- or alcohol-drinking mice infected with AAV-GFP or AAV-Rac1-DN. We observed a significant increase in the phosphorylation of LIMK and cofilin in AAV-GFP-infected animals during alcohol withdrawal (**Figure 2H-J**, **Table 1**) (Two-way ANOVA: pLIMK Effect of alcohol F_(1,20)_ = 35.12, p<0.0001; *Post hoc*: Water+AAV-GFP vs. Alcohol+AAV-GFP p<0.0001; pCofilin Effect of alcohol F_(1,20)_ = 380.5, p<0.0001; *Post hoc:* Water+AAV-GFP vs. Alcohol+AAV-GFP p<0.0001), which aligns with our prior results (**Figure 1K**). AAV-Rac1-DN overexpression had no effect on the phosphorylation of LIMK or cofilin in water-drinking animals (Two-way ANOVA: pLIMK Effect of virus F_(1,20)_ = 58.01, p<0.0001; *Post hoc*: Water+AAV-GFP vs. Water+AAV-Rac1-DN p = 0.3184; pCofilin Effect of virus F_(1,20)_ = 378.8, p<0.0001; *Post hoc:* Water+AAV-GFP vs. Water+AAV-Rac1-DN p = 0.9679). In contrast, overexpression of AAV-Rac1-DN significantly reduced alcohol-mediated phosphorylation of LIMK and cofilin (Two-way ANOVA: pLIMK Effect of virus x alcohol F_(1,20)_ = 105.1, p<0.0001; *Post hoc*: Alcohol+AAV-GFP vs. Alcohol+AAV-Rac1-DN p<0.0001; pCofilin Effect of virus x alcohol F_(1,20)_ = 423.8, p<0.0001; *Post hoc:* Alcohol+AAV-GFP vs. Alcohol+AAV-Rac1-DN p<0.0001). These results suggest that the molecular consequences of Rac1 stimulation by alcohol in the DMS is the phosphorylation, and therefore activation, of LIMK and the phosphorylation of its substrate cofilin.

### Alcohol promotes F-actin formation in the DMS via Rac1

We previously showed that chronic excessive alcohol intake increases F-actin assembly and decreases G-actin in the DMS (Laguesse et al., 2018). Phosphorylated cofilin is unable to cleave F-actin into G-actin (Bamburg, 1999). Therefore, we hypothesized that the consequence of the alcohol-mediated activation of the Rac1/LIMK/cofilin signaling is the formation of F-actin. To determine if actin remodeling in the DMS depends on Rac1 signaling, we examined the level of F-actin and G-actin in the DMS of mice that underwent 7 weeks of IA20%2BC or water and that were infected with AAV-GFP or AAV-Rac1-DN (**Figure 2F**, **Table 1**). Similar to previous findings (Laguesse et al., 2018), control mice infected with AAV-GFP in the DMS had a significantly higher level of F-actin after excessive chronic alcohol consumption in comparison to AAV-GFP-infected mice that drank water only (**Figure 2K-L**, **Table 1**) (Two-way ANOVA: Effect of alcohol F_(1,20)_ = 101.6, p<0.0001; *Post hoc*: Water+AAV-GFP vs. Alcohol+AAV-GFP p<0.0001). Conversely, mice infected with AAV-Rac1-DN had a significantly lower F-actin content in comparison to AAV-GFP mice both in the water- and alcohol-consuming groups (**Figure 2K-L**, **Table 1**) (Two-way ANOVA: Effect of virus F_(1,20)_ = 105.4, p<0.0001; *Post hoc*: Water+AAV-GFP vs. Water+AAV-Rac1-DN p = 0.0199; Alcohol+AAV-GFP vs. Alcohol+AAV-Rac1-DN p<0.0001). In addition, the magnitude of the difference in F-actin levels is greater between alcohol-drinking AAV-GFP and AAV-Rac1-DN mice compared to water-only animals (**Figure 2K-L**, **Table 1**) (Two-way ANOVA: Effect of virus x alcohol F_(1,20)_ = 30.92, p<0.0001). Overexpression of AAV-Rac1-DN had no effect on the level of G-actin in the water-only group (**Figure 2K**, **Figure 2M**). However, G-actin levels were reduced by alcohol in mice infected with AAV-GFP, which was reversed by overexpression of AAV-Rac1-DN (**Figure 2K**, **Figure 2M**, **Table 1**) (Two-way ANOVA: Effect of virus F_(1,20)_ = 7.585, p = 0.0122; Effect of alcohol F_(1,20)_ = 12.30, p = 0.0022; Effect of virus x alcohol F_(1,20)_ = 11.73, p = 0.0027; *Post hoc*: Water+AAV-GFP vs. Alcohol+AAV-GFP p = 0.0005; Water+AAV-Rac1-DN vs. Alcohol+AAV-GFP p = 0.0016; Alcohol+AAV-GFP vs. Alcohol+AAV-Rac1-DN p = 0.0018). Therefore, these data suggest that the alcohol-mediated increase in F-actin assembly in the DMS depends on Rac1.

### Rac1 promotes the remodeling of dendritic arbors in the DMS

F-actin is responsible for morphological remodeling in neurons (Honkura et al., 2008; Chazeau and Giannone, 2016; Costa et al., 2020). Previously, we showed that alcohol consumption increases dendritic branch complexity in DMS medium spiny neurons (MSNs) (Wang et al., 2015; Laguesse et al., 2018). Since alcohol leads to F-actin formation via Rac1, we hypothesized that the increase in dendritic branching after alcohol use is mediated via Rac1 signaling. To test this possibility, the DMS of mice that were subjected to IA20%2BC or water only was infected with a low titer of AAV-Rac1-DN or AAV-GFP (7×10^9^ - 3.5×10^10^ vg/ml). The goal was to infect a sparse population of neurons to allow for imaging of individual arbors (**Figure 3A**). Low-titer infection of Rac1-DN did not alter alcohol consumption (**Table 1**). Sholl analysis (Sholl, 1953) was then performed on the DMS of water- or alcohol-drinking mice infected with AAV-GFP or AAV-Rac1-DN (**Figure 3B-C**).

**Figure 3.**
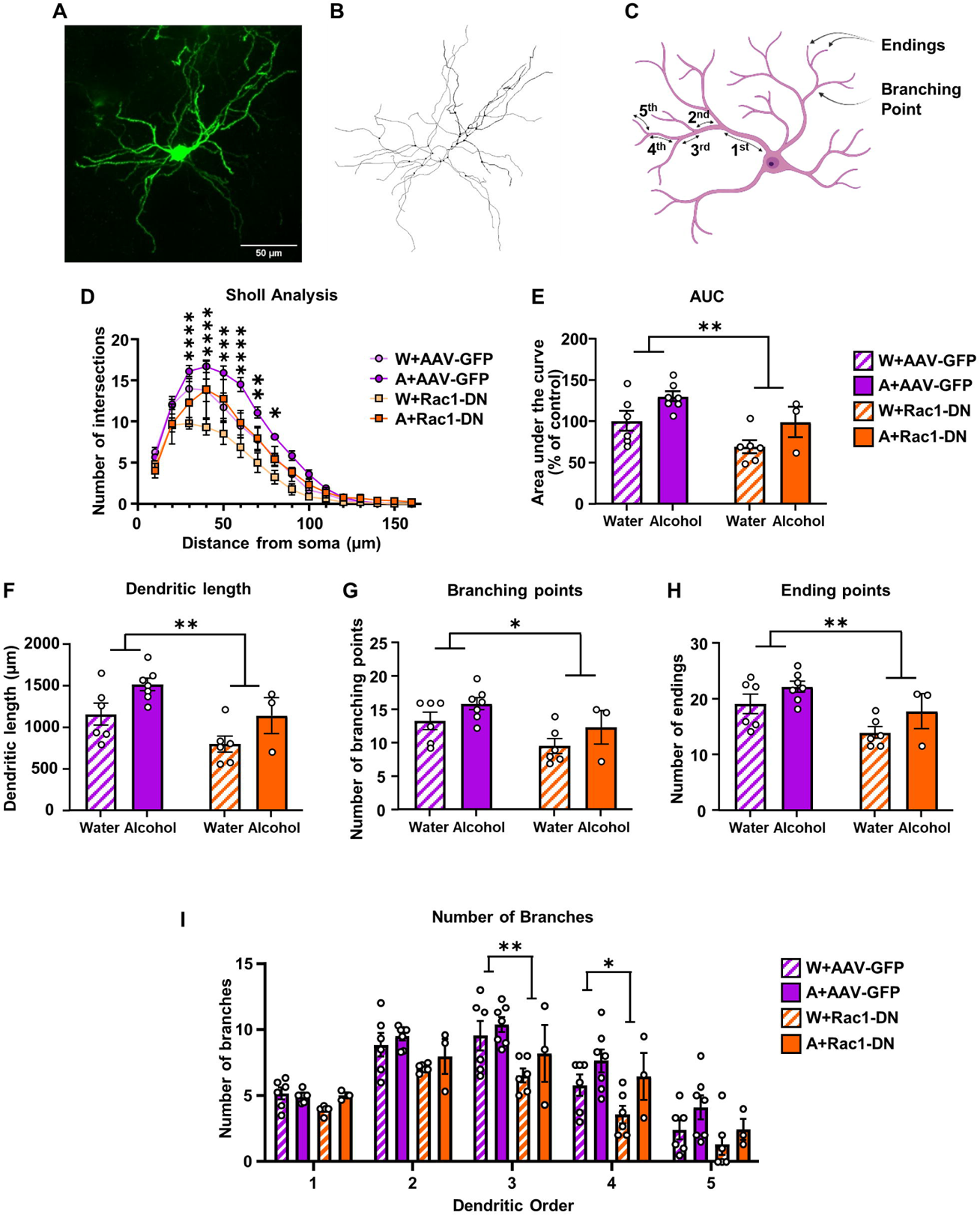
Rac1 promotes remodeling of dendritic arbors in the DMS. Low titer of AAV-GFP or AAV-Rac1-DN (7×10^9^ - 3.5×10^10^ vg/ml) was infused bilaterally into the DMS. After 1 week of recovery, mice underwent 7 weeks of IA20%2BC. Mice consuming water only were used as control. Twenty-four hours after the last drinking session, mice were perfused and brains were sliced into 100μm sections before MSN morphology was analyzed. **(A)** Sample image (40X magnification) of a GFP-positive DMS MSN. Scale bar, 50 μm. **(B)** Example of the reconstruction of the GFP-positive neuron. **(C)** Scheme of the morphological parameters measured in each neuron. Dendritic order increased after each branching point, defined as a dendritic intersection. **(D, E)** Analysis of neuronal dendritic arborization. Sholl analysis was performed on reconstructed neurons. The number of intersections was analyzed for each point **(D)** and the area under the curve (AUC) **(E)** was calculated. **(F)** Measurement of total dendritic length. **(G)** Number of branching points. **(H)** Number of ending points. **(I)** Number of branches per dendritic order. Data are represented as mean ± SEM and analyzed by two-way ANOVA. Main effect of virus is represented graphically. \**p* < 0.05, \*\**p*<0.01, \*\*\**p* < 0.001. Water+AAV-GFP: 19 neurons from 6 mice; Alcohol+AAV-GFP: 23 neurons from 7 mice; Water+AAV-Rac1-DN: 15 neurons from 6 mice, Alcohol+AAV-Rac1-DN: 10 neurons from 3 mice.

Similar to our prior findings (Wang et al., 2015; Laguesse et al., 2018), we found that alcohol significantly increased the complexity of dendritic trees of DMS MSNs (**Figure 3D**) (Three-way ANOVA: Effect of alcohol x distance F_(15,270)_ = 5.227, p < 0.0001). Specifically, DMS dendrites 30-80μm from the soma exhibited significantly more intersections in alcohol-drinking vs. water-only-drinking mice (**Figure 3D**) (Two-way ANOVA: Effect of alcohol F_(1, 20)_ = 9.063, p < 0.0001, *Post hoc Šídák’s multiple comparisons test Water vs. Alcohol*). The increase in dendritic complexity was further quantified by calculating the area under the curve (AUC) (**Figure 3E**) (Two-way ANOVA: Effect of alcohol F_(1, 18)_ = 7.932, p = 0.0114). In addition, we observed that dendritic length (**Figure 3F**), the number of branching points (**Figure 3G**), and the number of ending points (**Figure 3H**), were also increased by alcohol (Two-way ANOVA: Effect of alcohol Dendritic Length F_(1, 18)_ = 8.247, p = 0.0101; Branching points F_(1, 18)_ = 4.050, p = 0.0594; Ending points F_(1,18)_ = 4.759, p = 0.0426).

In contrast, infection of the DMS with AAV-Rac1-DN significantly reduced dendritic arborization (**Figure 3D**) (Three-way ANOVA: Effect of virus x distance F_(15, 270)_ = 4.890, p < 0.0001). Specifically, the number of dendritic intersections 30-80μm from the soma was significantly decreased when DMS neurons were infected with AAV-Rac1-DN as compared to neurons infected with AAV-GFP (**Figure 3D**) (Two-way ANOVA of consolidated data: Effect of virus F_(1, 20)_ = 10.71, p = 0.0038, *Post hoc Šídák’s multiple comparisons test, AAV-GFP vs. AAV-Rac1-DN*). Furthermore, AAV-Rac1-DN infection significantly decreased AUC (**Figure 3E**), along with dendritic length, branching points, and ending points (**Figure 3F-H**) (Two-way ANOVA: Effect of virus AUC F_(1, 18)_ = 8.754, p = 0.0084, Dendritic length F_(1, 18)_ = 9.186, p = 0.0072, Branching points F_(1, 18)_ = 7.551, p = 0.0132, Ending points F_(1, 18)_ = 9.030, p = 0.0076). However, there was no interaction between the variables (Three-way ANOVA: Effect of alcohol x virus x distance F_(15,270)_ = 0.2661, p = 0.9975; Two-way ANOVA: Effect of alcohol x virus AUC F_(1,18)_ = 0.0005846, p = 0.9810, Dendritic length F_(1,18)_ = 0.004819, p = 0.9454, Branching points F_(1,18)_ = 0.005694, p = 0.9407, Ending points F_(1,18)_ = 0.06334, p = 0.8041). Next, we examined the number of branches per dendritic order (**Figure 3I**). Three-way ANOVA showed a significant main effect of virus (F_(1,18)_ = 7.859, p = 0.0117) and a significant main effect of alcohol (F_(1,18)_ = 4.921, p = 0.0396) but no interaction between virus and alcohol (F_(1,18)_ = 0.2476, p = 0.6248). Next, the number of branches was analyzed for each order. Two-way ANOVA showed a significant main effect of virus at 3^rd^ and 4^th^ order, and a significant main effect of alcohol at 4^th^ order (**Figure 3I**).

Together, these data confirmed that chronic excessive alcohol use significantly increases the complexity of dendritic branching in DMS MSNs (Wang et al., 2015; Laguesse et al., 2018). Importantly, we show that Rac1 contributes to dendritic branching in DMS neurons.

### Rac1 in the DMS is required for the alcohol-mediated alteration in dendritic spine morphology

Actin is the major structural protein in the post-synaptic density (PSD) (Ratner and Mahler, 1983). Actin cytoskeleton organization directly controls dendritic spine morphology (Schubert and Dotti, 2007; Hotulainen and Hoogenraad, 2010; Basu and Lamprecht, 2018). Rac1 signaling controls actin dynamics and is involved in dendritic spine morphogenesis (Bosco et al., 2009; Haditsch et al., 2009; Haditsch et al., 2013; Costa et al., 2020). Since Rac1 is activated by alcohol which in turn promotes F-actin assembly we hypothesized that the dendritic spines in the DMS are altered by alcohol in a Rac1-dependent manner.

Next, we examined the shape of individual dendritic spines on distal dendritic branches (3^rd^ or 4^th^ order) in mice infected with AAV-GFP and AAV-Rac1-DN in DMS and that consumed 20% alcohol or water only for 7 weeks (**Figure 4A**). Spine density (spine number per 10µm) was unaffected by alcohol drinking or AAV-Rac1-DN infection (**Figure 4B**) (Two-way ANOVA: Effect of virus F_(1,15)_ = 0.1118, p = 0.7428; Effect of alcohol F_(1,15)_ = .01518, p = 0.9036; Effect of virus x alcohol F_(1,15)_ = 1.150, p = 0.3006).

**Figure 4.**
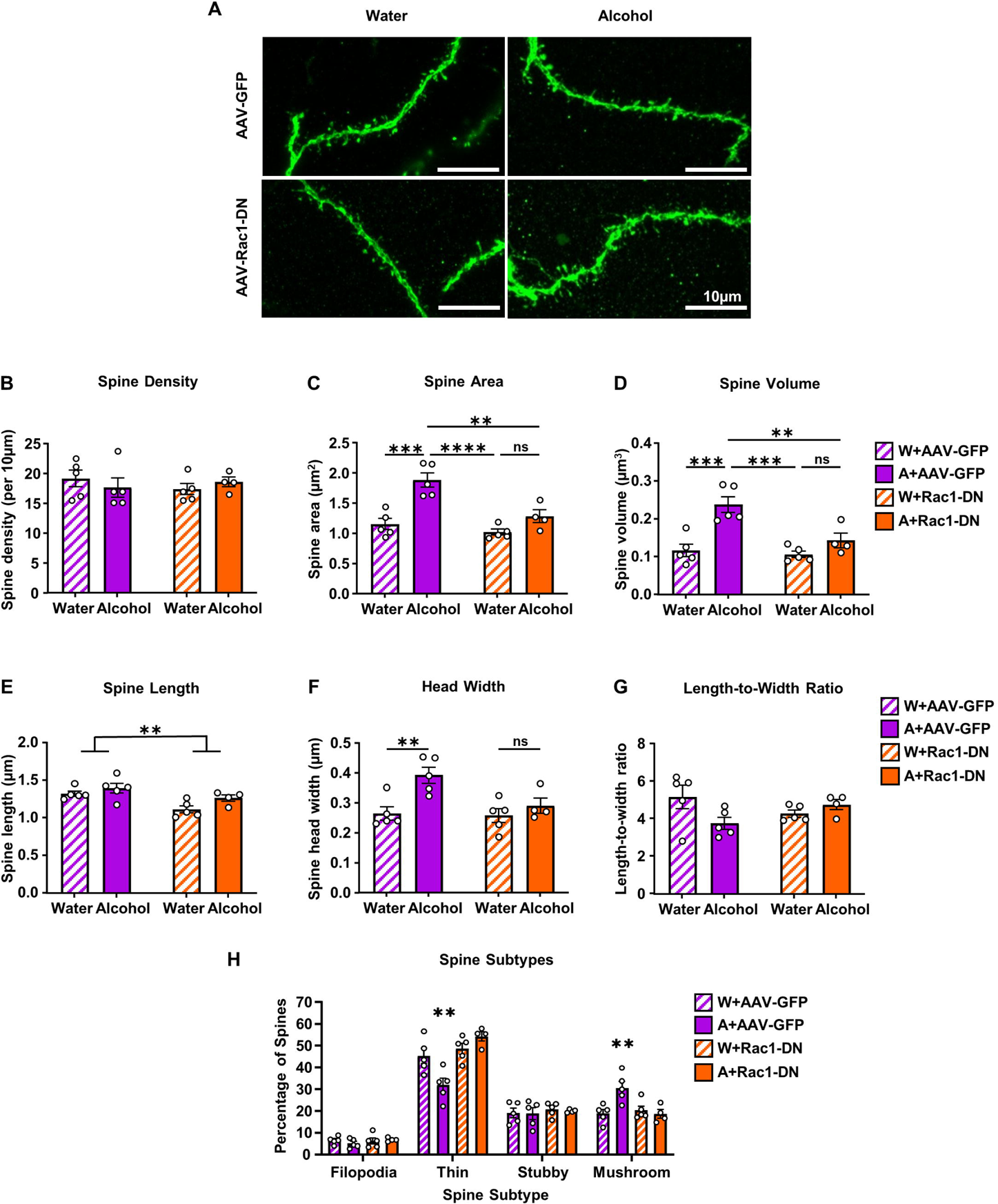
Rac1 in the DMS is required for the alcohol-mediated morphological changes in dendritic spines. Low titer of AAV-GFP or AAV-Rac1-DN (7×10^9^ - 3.5×10^10^ vg/ml) was infused bilaterally into the DMS. After 1 week of recovery, mice underwent 7 weeks of IA20%-2BC. Mice consuming water only were used as control. Twenty-four hours after the last drinking session, mice were perfused, and MSN dendritic spine morphology was analyzed. **(A)** Sample image of a GFP-positive DMS MSN (100X magnification) from each group. Scale bar, 10 μm. **(B)** Average dendritic spine density (number of spines/10 µm). **(C)** Average dendritic spine area (µm^2^). **(D)** Average dendritic spine volume (µm^3^). **(E)** Average dendritic spine length (µm). **(F)** Average diameter of dendritic spine heads (µm). **(G)** Dendritic spine length-to-width ratio. Data is represented as mean ± SEM and analyzed using two-way ANOVA and Šidák’s *post hoc* multiple comparisons test. *p<0.05, **p<0.01, ***p<0.001. ****p<0.0001. **(H)** Percentage of filopodia, thin, stubby, and mushroom dendritic spines for water- and alcohol-drinking mice infected with either AAV-GFP or AAV-Rac1-DN. Three-way ANOVA and Šidák’s *post hoc* multiple comparisons test. ****p<0.0001. Water+AAV-GFP: 14 neurons from 5 mice; Alcohol+AAV-GFP: 13 neurons from 5 mice; Water+AAV-Rac1-DN: 14 neurons from 5 mice; Alcohol+AAV-Rac1-DN: 11 neurons from 4 mice.

We found that in control mice infected with AAV-GFP, alcohol drinking significantly increased dendritic spine area, aligning with prior findings (Laguesse et al., 2018) (Two-way ANOVA: Effect of alcohol F_(1,15)_ = 27.17, p = 0.0001) (**Figure 4C**). Alcohol drinking also significantly increased dendritic spine volume (Two-way ANOVA: Effect of alcohol F_(1,15)_ = 21.79, p = 0.0003) (**Figure 4D**). Both spine length (**Figure 4E**) and average spine head width (**Figure 4F**) were significantly increased by alcohol drinking (Two-way ANOVA: Spine Length Effect of alcohol F_(1,15)_ = 4.617, p = 0.0484; Head Width Effect of alcohol F_(1,15)_ = 10.69, p = 0.0052).

In contrast, the alcohol-mediated alterations in dendritic spine morphology were attenuated by Rac1 inhibition. Specifically, Rac1 inhibition significantly decreased dendritic spine area in alcohol-drinking animals, without affecting water-only controls. (**Figure 4C**) (Two-way ANOVA: Effect of virus F_(1,15)_ = 14.86, p = 0.0016; Effect of virus x alcohol F_(1,15)_ = 6.075, p = 0.0263; *Post hoc Šídák’s multiple comparisons test: W+AAV-GFP vs. A+AAV-GFP p = 0.0003, A+AAV-GFP vs. W+Rac1-DN p < 0.0001, A+AAV-GFP vs. A+AAV-Rac1-DN p = 0.0029, W+AAV-Rac1-DN vs. A+AAV-Rac1-DN p = 0.2739*). The alcohol-mediated increase in dendritic spine volume was also prevented by Rac1-DN, without affecting water-only controls (**Figure 4D**) (Two-way ANOVA: Effect of virus F_(1,15)_ = 9.634, p = 0.0073; Effect of virus x alcohol F_(1,15)_ = 6.172, p = 0.0253; *Post hoc Šídák’s multiple comparisons test: W+AAV-GFP vs. A+AAV-GFP p = 0.0005, A+AAV-GFP vs. W+Rac1-DN p = 0.0002, A+AAV-GFP vs. A+AAV-Rac1-DN p = 0.0078, W+AAV-Rac1-DN vs. A+AAV-Rac1-DN p = 0.4614*). In addition, we found that AAV-Rac1-DN slightly reduced spine length independent of drinking (**Figure 4E**), and decreased spine head width (**Figure 4F**) (Two-way ANOVA: Spine Length Effect of virus F_(1,15)_ = 10.99, p = 0.0047; Effect of virus x alcohol F_(1,15)_ = 0.5682, p = 0.4626; Head Width Effect of virus F_(1,15)_ = 5.064, p = 0.0399; Effect of virus x alcohol F_(1,15)_ = 3.728, p = 0.0726; *Post hoc Šídák’s multiple comparisons test: W+AAV-GFP vs. A+AAV-GFP p = 0.0086, W+AAV-Rac1-DN vs. A+AAV-Rac1-DN p = 0.7946*). Length-to-width ratio was not affected by alcohol or Rac1 inhibition alone, but a combination of these factors (**Figure 4G**) (Two-way ANOVA: Effect of alcohol F_(1,15)_ = 1.285, p = 0.2748; Effect of virus F_(1,15)_ = 0.0.01974, p = 0.8901; Effect of virus x alcohol F_(1,15)_ = 5.418, p = 0.0343). Together, these findings suggest that Rac1 contributes to the alcohol-mediated increases in dendritic spine area, volume, and length-to-width ratio, while also directly affecting spine length and head width.

We then examined whether alcohol-mediated activation of Rac1 signaling and the increase in F-actin content was responsible for the maturation of DMS MSN dendritic spines. Dendritic spines are classed into four subtypes: filipodia, thin, stubby, and mushroom, with filopodia being the least mature form of spines and mushrooms being the most mature form (Kasai et al., 2003). Spines are characterized by different shape, with primary variables being spine length, neck, and head width (Kasai et al., 2003). Like our previous study (Laguesse et al., 2018), we found that alcohol drinking increased the maturity of dendritic spines in the DMS. Specifically, we found that in AAV-GFP infected mice, alcohol significantly increased the proportion of mushroom-shaped spines at the expense of thin spines, while not affecting filopodia or stubby spines (**Figure 4H**). Rac1 inhibition blocked the alcohol-mediated increase in mushroom spines and decrease in thin spines but had no effect on animals drinking water only (**Figure 4H**) (Three-way ANOVA: Effect of virus F_(1, 15)_ = 2.745, p = 0.1183; Effect of alcohol F_(1, 15)_ = 0.003263, p = 0.9552; Spine type F_(3, 45)_ = 304.5, p < 0.0001; Effect of virus x spine type F_(3, 45)_ = 16.83, p < 0.0001; Effect of alcohol x spine type F_(3, 45)_ = 4.052, p = 0.0124, Effect of virus x alcohol x spine type F_(3, 45)_ = 13.08, p < 0.0001; *Post hoc Šídák’s multiple comparisons test: Thin: W+AAV-GFP vs. A+AAV-GFP p = 0.0018, A+AAV-GFP vs. W+AAV-Rac1-DN p < 0.0001, A+AAV-GFP vs. A+AAV-Rac1-DN p < 0.0001; Mushroom: W+AAV-GFP vs. A+AAV-GFP p = 0.0082, A+AAV-GFP vs. W+AAV-Rac1-DN p = 0.0304, A+AAV-GFP vs. A+AAV-Rac1-DN p = 0.0123*). Together, our data suggest that Rac1 signaling is responsible for the alcohol-mediated morphological changes in dendritic spine structure and maturation in DMS MSNs.

### Rac1 does not play a role in the development and maintenance of voluntary alcohol intake

Next, we set to determine if the alcohol-dependent, Rac1-mediated molecular and cellular changes have consequences on alcohol-related behaviors. First, we tested the effect of Rac1 inhibition on the development of excessive alcohol drinking. To do so, the DMS of mice was bilaterally infused with AAV-Rac1-DN or AAV-GFP (3.5×10^12^ vg/ml). After allowing for three weeks of recovery, animals underwent IA20%2BC for 7 weeks and drinking and preference were measured (**Figure 5A**). We found that there was no difference in alcohol consumption in the group with overexpressed AAV-Rac1-DN in the DMS compared to AAV-GFP controls (**Figure 5B**) (RM Two-way ANOVA: Effect of virus F_(1,17)_ = 0.08351, p = 0.7761; Effect of session F_(6, 102)_ = 5.420, p < 0.0001; Effect of virus x session F_(6, 102)_ = 0.1339, p = 0.9916). Water drinking was also unaltered (**Figure 5C**). RM Two-way ANOVA: Effect of virus F_(1,17)_ = 0.4571, p = 0.5081; Effect of session F_(6,102)_ = 3.855, p = 0.0017; Effect of virus x session F_(6,102)_ = 0.8631, p = 0.5248). As a result, alcohol preference was also unchanged (**Figure 5D**) (RM Two-way ANOVA: Effect of virus F_(1,17)_ = 0.1798, p = 0.6768; Effect of session F_(6, 102)_ = 3.196, p = 0.0065; Effect of virus x session F_(6, 102)_ = 0.4250, p = 0.8608).

**Figure 5.**
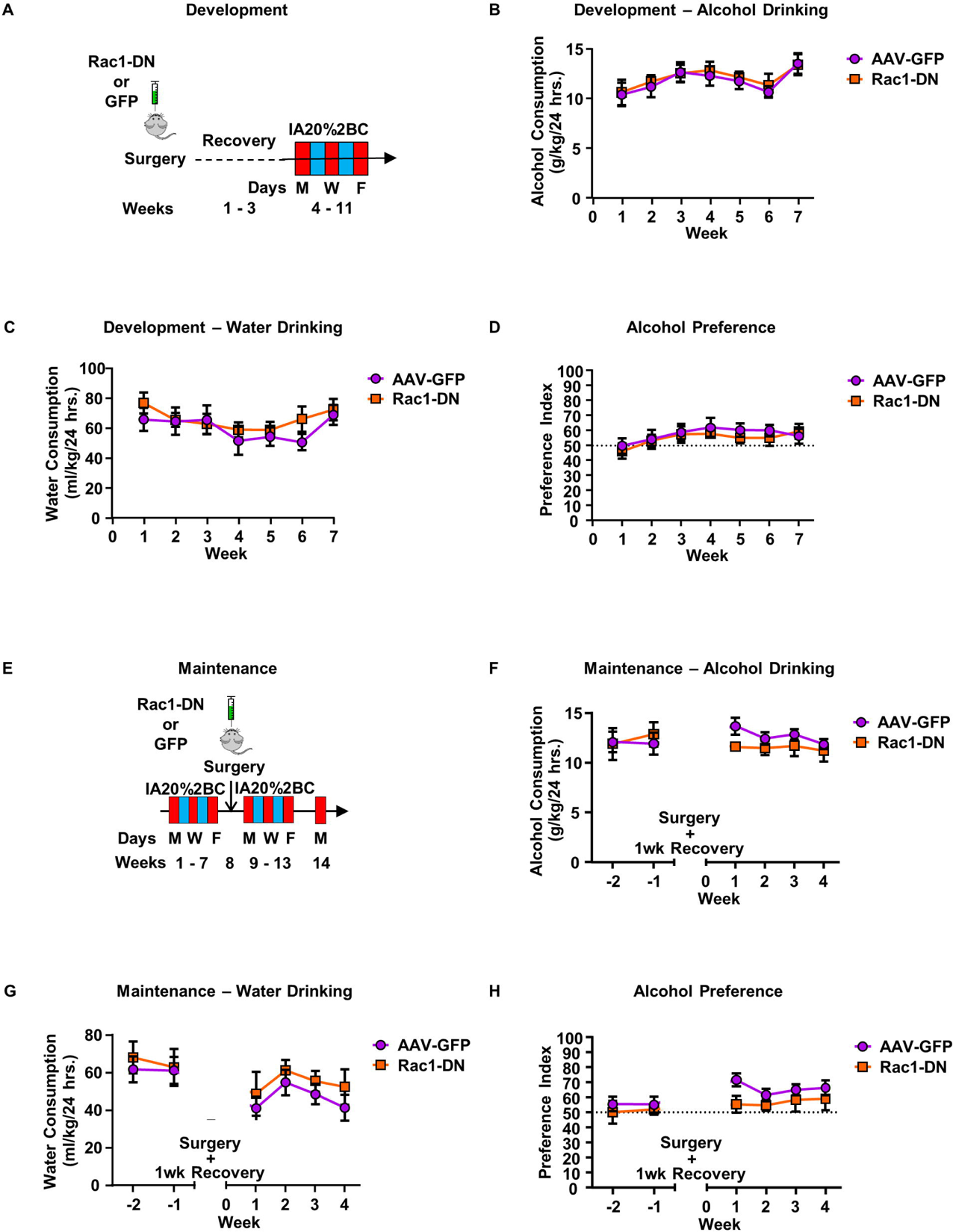
Rac1 does not play a role in the development and maintenance of voluntary alcohol intake. **(A)** Development: Surgery and IA20%2BC timeline. Mice received a bilateral, dual-infusion of AAV-Rac1-DN or AAV-GFP as a control. After 3 weeks of recovery, IA20%2BC began and continued for 7 weeks. **(B)** Weekly average alcohol consumption (g/kg/24hrs.) of development drinking mice. **(C)** Water consumption averages for the IA20%2BC development drinking groups. **(D)** Preference for alcohol during the development IA20%2BC drinking, with an index above 50 indicating a preference for alcohol. **(E)** Maintenance: Surgery and IA20%2BC timeline. Mice first underwent IA20%2BC for 7 weeks. Groups were balanced based on average daily alcohol consumption in last 6 drinking sessions. Mice received a bilateral, dual-infusion of AAV-Rac1-DN or AAV-GFP as a control and were allowed one full week for recovery. IA20%2BC resumed for 4 weeks. **(F)** Weekly average consumption (g/kg/24hrs.) for maintenance drinking mice. **(G)** Water consumption average for maintenance drinking mice. **(H)** Preference for alcohol for maintenance-drinking mice. All data represented as mean ± SEM and analyzed with two-way ANOVA with repeated measures. n = 9-10.

Next, we determined the potential contribution of Rac1 to the maintenance of alcohol-drinking behavior. Mice were first subjected to the IA20%2BC paradigm for 7 weeks. Experimental and control groups were balanced based on drinking average. Mice underwent surgery to overexpress AAV-Rac1-DN, or AAV-GFP as a control, in the DMS. IA20%2BC was then resumed for 4 weeks and alcohol drinking was evaluated (**Figure 5E**). Overexpression of AAV-Rac1-DN did not alter the maintenance of alcohol drinking (**Figure 5F**) (RM Two-way ANOVA: Effect of virus F_(1,14)_ = 2.454, p = 0.1396; Effect of session F_(3,42)_ = 1.262, p = 0.2998; Effect of virus x session F_(3,42)_ = 0.5923, p = 0.6235). Water consumption was similarly unchanged (**Figure 5G**) (RM Two-way ANOVA: Effect of virus F_(1,13)_ = 0.8287, p = 0.3792; Effect of session F_(3, 39)_ = 3.606, p = 0.0216; Effect of virus x session F_(3,39)_ = 0.1149, p = 0.9508), leading to similar alcohol preference between the group (**Figure 5H**) (RM Two-way ANOVA: Effect of virus F_(1, 14)_ = 2.520, p = 0.1347; Effect of session F_(3,42)_ = 0.7839, p = 0.5096; Effect of virus x session F_(3,42)_ = 0.7518, p = 0.5275). Together, these results suggest that Rac1 in the DMS does not contribute to the development or maintenance of voluntary alcohol drinking in the IA20%2BC paradigm.

### Rac1 in the DMS is required for alcohol-associated goal-directed learning

We observed that the activation of Rac1 signaling after chronic excessive alcohol use was specific to the DMS, a brain region that is associated with goal-directed learning (Dolan and Dayan, 2013; Shan et al., 2014). As goal-directed behavior is particularly important for drug seeking (Singer et al., 2018), we hypothesized that Rac1 in the DMS may play a role in alcohol-associated goal-directed learning.

To examine this hypothesis, the DMS of mice was bilaterally infused with AAV-Rac1-DN or AAV-GFP (3.5×10^12^ vg/ml). Mice were allowed 3 weeks to recover before undergoing IA20%2BC for 7 weeks. Mice were then trained to self-administer 20% alcohol on a fixed ratio of reinforcement 1 (FR1) schedule for 20 hours over four sessions during the first week before transitioning to 2-hour sessions. After eight 2-hour sessions of FR1, mice progressed to FR2, and alcohol-associated active lever presses and inactive lever presses were examined (**Figure 6A**). We found that there was no significant change in active lever presses between mice infected with AAV-Rac1-DN or AAV-GFP (**Figure 6B**, **Table 1**) (RM Two-way ANOVA: Effect of virus F_(13, 143)_ = 1.522, p = 0.1163; Effect of session F_(3.043, 33.47)_ = 1.232, p = 0.3137; Effect of virus x session F_(13, 143)_ = 1.522, p = 0.1163). However, mice infected with AAV-Rac1-DN pressed the inactive lever significantly more than the AAV-GFP-infected mice (**Figure 6C**) (RM Two-way ANOVA: Effect of virus F_(1, 11)_ = 5.102, p = 0.0452; Effect of session F_(13, 143)_ = 0.9024, p = 0.5522; Effect of virus x session F_(13, 143)_ = 0.8569, p = 0.5996). This phenotype is exemplified in the discrimination index, showing that the AAV-Rac1-DN mice exhibited significantly worse discrimination for the active lever throughout the OSA experiment in comparison to AAV-GFP controls (**Figure 6D**) (RM Two-way ANOVA: Effect of virus F_(1, 11)_ = 5.320, p = 0.0415; Effect of session F_(13, 143)_ = 2.237, p = 0.0107; Effect of virus x session F_(13,143)_ = 0.8674, p = 0.5886). A similar phenotype was observed in the proportion of rewarded lever presses, which accounts for active lever presses that do not lead to a reward. Specifically, while over 50% of presses of the AAV-GFP control animals led to a reward, the AAV-Rac1-DN group failed to reach this threshold throughout the testing period (**Figure 6E**) (RM Two-way ANOVA: Effect of virus F_(1,11)_ = 16.66, p = 0.0018, Effect of session F_(13,143)_ = 6.400, p < 0.0001; Effect of virus x session F_(13,143)_ = 0.5518, p = 0.8880). There was no difference in alcohol consumption between groups (**Figure 6F**) (RM Two-way ANOVA: Effect of virus F_(1,11)_ = 1.099, p = 0.3169; Effect of session F_(13, 143)_ = 15.18, p < 0.0001; Effect of virus x session F_(13, 143)_ = 1.156, p = 0.3178), which aligns with the IA20%2BC data (**Figure 5**). Examination of the inter-response interval between lever presses of the final session shows that AAV-Rac1-DN mice exhibited a significantly increased proportion of lever presses in short intervals, and significantly less lever presses in the longer intervals, compared to AAV-GFP controls (**Figure 6G**) (RM Two-way ANOVA: Effect of virus F_(1,11)_ = 0.5360, p = 0.4794; Effect of interval F_(4,44)_ = 64.48, p < 0.0001; Effect of virus x interval F_(4, 44)_ = 6.145, p = 0.0005; *Post hoc* 0-5 p = 0.0034, >20 p < 0.001). These phenotypes can be visualized in the behavioral trace of individual mouse profiles from the OSA session with the greatest discrimination difference (**Figure 6H**). As the DMS plays an important role in movement (Kravitz and Kreitzer, 2012), we examined the consequence of Rac1 inhibition in the DMS on locomotion using an open field paradigm. As shown in **Figure 6l**, attenuation of Rac1 activity in the DMS had no effect on locomotion (Unpaired t-test: *t_(_*_16)_ = 0.06783, p = 0.9468). Therefore, this behavioral difference is not due to changes in motor behavior. Together, these data suggest that Rac1 in the DMS plays a role in discrimination of alcohol-associated rewarded lever presses.

**Figure 6.**
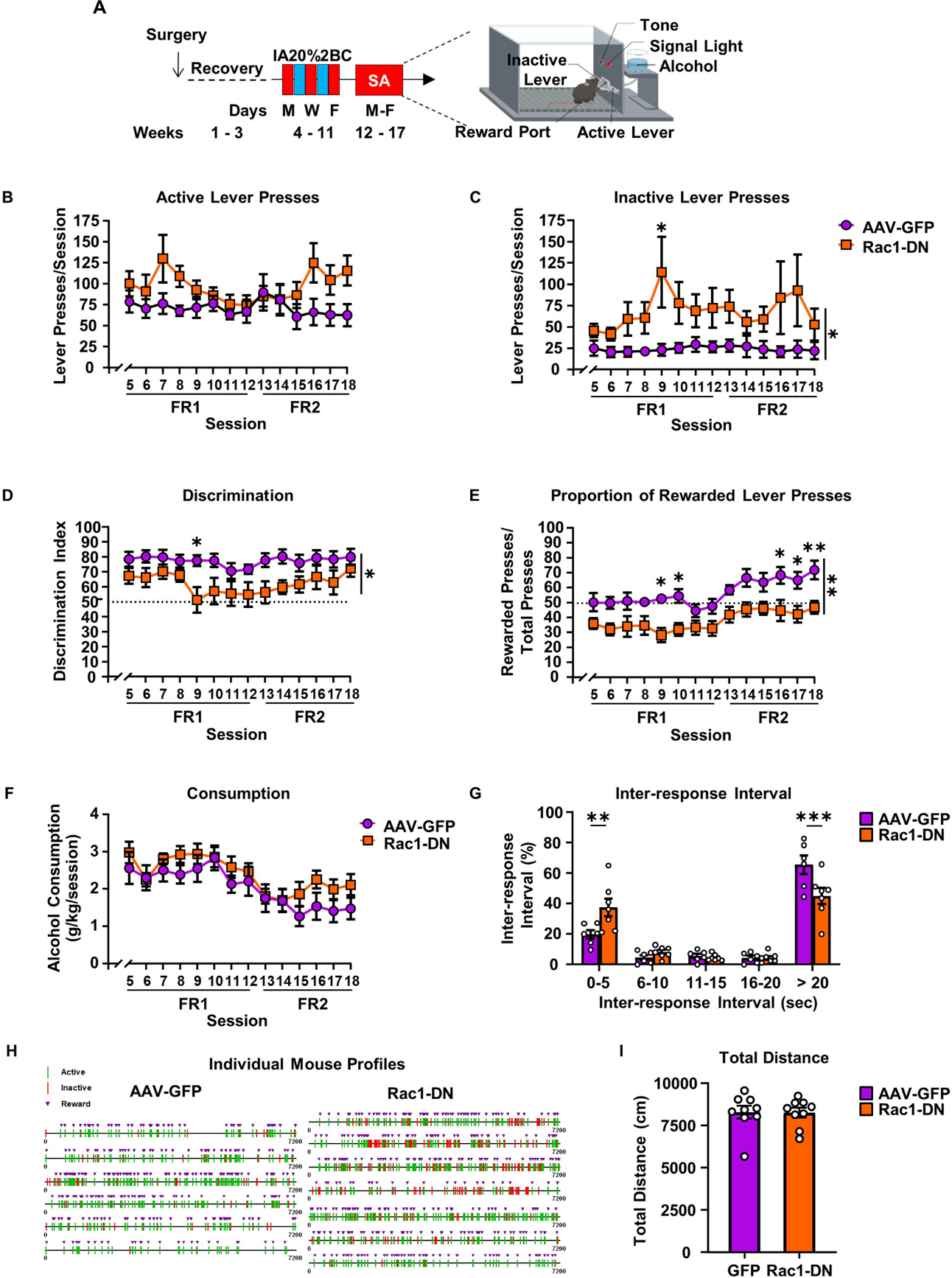
Rac1 in the DMS is required for alcohol-associated goal-directed learning. **(A)** Operant self-administration of 20% alcohol paradigm. Mice underwent surgery to overexpress AAV-Rac1-DN or AAV-GFP in the DMS. After 3 weeks of recovery, mice underwent 7 weeks of IA20%2BC to develop a preference for alcohol before transitioning to 20% alcohol OSA for 1 week of training and 4 experimental weeks. **(B-C)** The group average of active **(B)** and inactive **(C)** lever presses during the 2-hour sessions. **(D)** Discrimination index for the active lever. Calculated as proportion of active lever presses compared to total lever presses. **(E)** Proportion of rewarded lever presses. Calculated as rewarded lever presses compared to total lever presses. **(F)** Alcohol consumption during each session. **(G)** Proportion of inter-response interval between lever presses during the final session. Data are presented as mean ± SEM and analyzed using RM Two-way ANOVA and Šidák’s *post hoc* multiple comparisons test. *p<0.05, **p<0.01, ***p<0.001. n = 6, 7. **(H)** Individual mouse behavioral profiles from OSA session 9. Each line corresponds with an active (green) or inactive (red) lever press, and triangles represent reward (purple). **(I)** Total distance moved in an open field. Mice were placed in an open field and locomotion was recorded for 20 minutes. Data represented as mean ± SEM and analyzed using an unpaired t-test. n = 9 per group.

Next, we examined if the reduction in discrimination as a result of Rac1-DN is related to deficits in goal-directed behavior. Goal-directed behavior is reliant on an association between response and outcome (Yin et al., 2005; Balleine and O’Doherty, 2010). We used a contingency degradation model to test the hypothesis that Rac1 in the DMS is involved in alcohol-associated goal-directed behavior. Mice were first infected with AAV-Rac1-DN or AAV-GFP in the DMS before undergoing IA20%2BC for 7 weeks. Mice were then trained to operant self-administer alcohol as described in (Morisot et al., 2019a) (**Figure 7A**). To bias mice towards goal-directed behavior, mice were trained on a random ratio (RR) schedule of reinforcement, which is known to promote goal-directed actions (Yin et al., 2005) (**Figure 7A**). AAV-GFP-infected mice significantly reduced active lever pressing during degraded sessions compared to non-degraded sessions (**Figure 7B**) However, AAV-Rac1-DN-infected mice pressed similarly during non-degraded and degraded sessions indicating a disruption in alcohol-associated, goal-directed learning (**Figure 7B**) (Two-way ANOVA: Effect of virus F_(1,12)_ = 4.863, p = 0.0477; Effect of degradation F_(1,12)_ = 17.98, p = 0.0011; Effect of virus x degradation F_(1,12)_ = 11.38, p = 0.0055; *Post hoc* ND vs. D: AAV-GFP p = 0.0003, AAV-Rac1-DN p = 0.7989). As shown in **Figure 7C**, the port entries during non-degraded and degraded sessions were unchanged (Two-way ANOVA: Effect of virus F_(1,2_ = 1.060, p = 0.3236; Effect of degradation F_(1,12)_ = 4.732, p = 0.0503; Effect of virus x degradation F_(1,12)_ = 0.4249, p = 0.5268). Together, these data suggest that Rac1 in the DMS plays a role in alcohol-associated, goal-directed learning.

**Figure 7.**
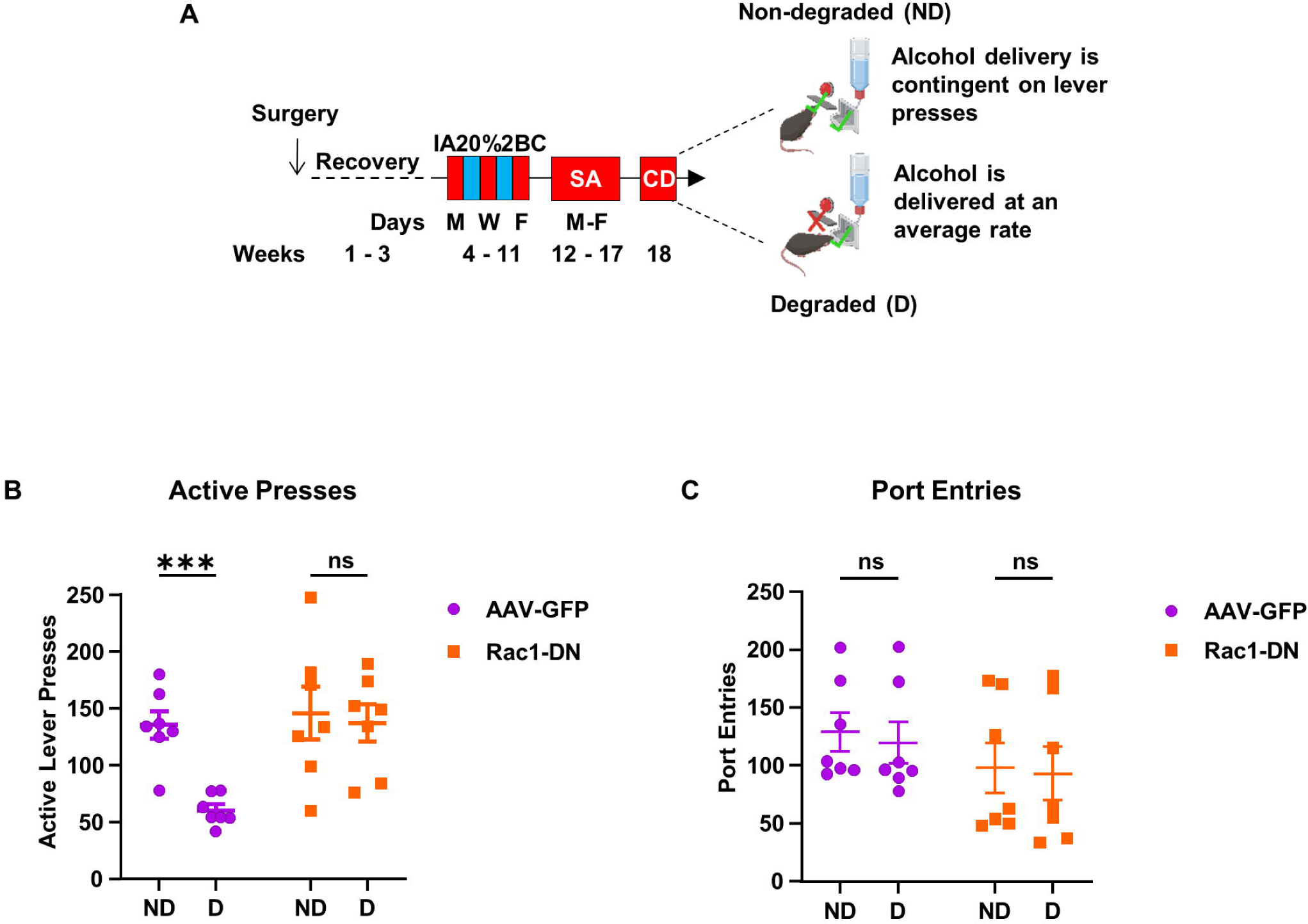
Rac1 in the DMS is required for alcohol action-outcome associations. **(A)** Contingency degradation paradigm. Mice received a dual-infusion of AAV-GFP or AAV-Rac1-DN into the DMS. After a recovery period, mice underwent IA20%2BC for 7 weeks before being trained to operant self-administer alcohol. Mice were first trained on an FR1 schedule before progressing to a random ratio (RR) schedule of reinforcement. Under RR, mice received a reward after a random number of active lever presses within a range (e.g. RR2 1-3 presses, RR3 2-4 presses, RR4 3-5 presses). Mice were trained for 5 sessions at each reinforcement step and spent 10 sessions at RR4 before testing. During the contingency degradation test, there were two types of sessions: non-degraded (ND) and degraded (D). During non-degraded sessions, rewards were delivered on an RR4 schedule. During degraded sessions, rewards were delivered at a rate equal to the average of the last week of training, but active lever presses had no effect. Testing consisted of a ND session followed by a D session, repeated three times. **(B)** Average active lever presses during ND and D testing sessions for AAV-GFP and AAV-Rac1-DN mice. **(C)** Average port entries during ND and D testing sessions for AAV-GFP and AAV-Rac1-DN mice. Data represented as mean ± SEM and analyzed using two-way ANOVA with Šidák’s *post hoc* multiple comparisons test. ***p<0.001. n = 7 per group.

### Rac1 in the DMS is not required for sucrose goal-directed learning

Alcohol is often distinct from natural reward (Alhadeff et al., 2019; Nall et al., 2021; Martins et al., 2022), and Rac1 in the DMS is not activated in response to sucrose consumption (**Figure 1I**). We hypothesized that Rac1 in the DMS does not play a role in sucrose-dependent goal-directed learning. To test this hypothesis, the DMS of mice was first bilaterally infused with AAV-Rac1-DN or AAV-GFP (3.5×10^12^ vg/ml). Mice underwent IA20%2BC for 7 weeks to match the condition of alcohol self-administration. The mice were then trained to operant self-administer 1% sucrose, initially starting with an FR1 reinforcement schedule for 20 hours of training over four sessions before transitioning to 2-hour sessions. After 8 2-hour sessions of FR1, mice were advanced to an FR2 schedule (**Figure 8A**). We observed no difference between mice with AAV-Rac1-DN and AAV-GFP overexpression in active lever pressing (**Figure 8B**), inactive lever pressing (**Figure 8C**), discrimination (**Figure 8D**), or proportion of rewarded lever presses (**Figure 8E**) (RM Two-way ANOVA: **8B** Effect of virus F_(1,11)_ = 4.651, p = 0.0540; Effect of session F_(3.905, 42.36)_ = 5.335, p = 0.0015; Effect of virus x session F_(13,141)_ = 0.9458, p = 0.5079; **8C** Effect of virus F_(1,11)_ = 0.2087, p = 0.6566; Effect of session F_(2.224, 24.12)_ = 0.8833, p = 0.4364; Effect of virus x session F_(13,141)_ = 0.6457, p = 0.8119; **8D** Effect of virus F_(1,11)_ = 0.007081, p = 0.9344; Effect of session F_(3.161, 34.29)_ = 1.888, p = 0.1475; Effect of virus x session F_(13,141)_ = 0.7552, p = 0.7057; **8E** Effect of virus F_(1,11)_ = 0.0001974, p = 0.9890; Effect of session F_(3.512, 38.10)_ = 6.534, p = 0.0007; Effect of virus x session F_(13,141)_ = 0.7314, p = 0.7298). These data suggest that Rac1 in the DMS does not affect learning of natural-reward-associated goal-directed behavior and that the learning deficiency exhibited by AAV-Rac1-DN overexpression in the DMS is specific to alcohol-associated goal-directed learning.

**Figure 8.**
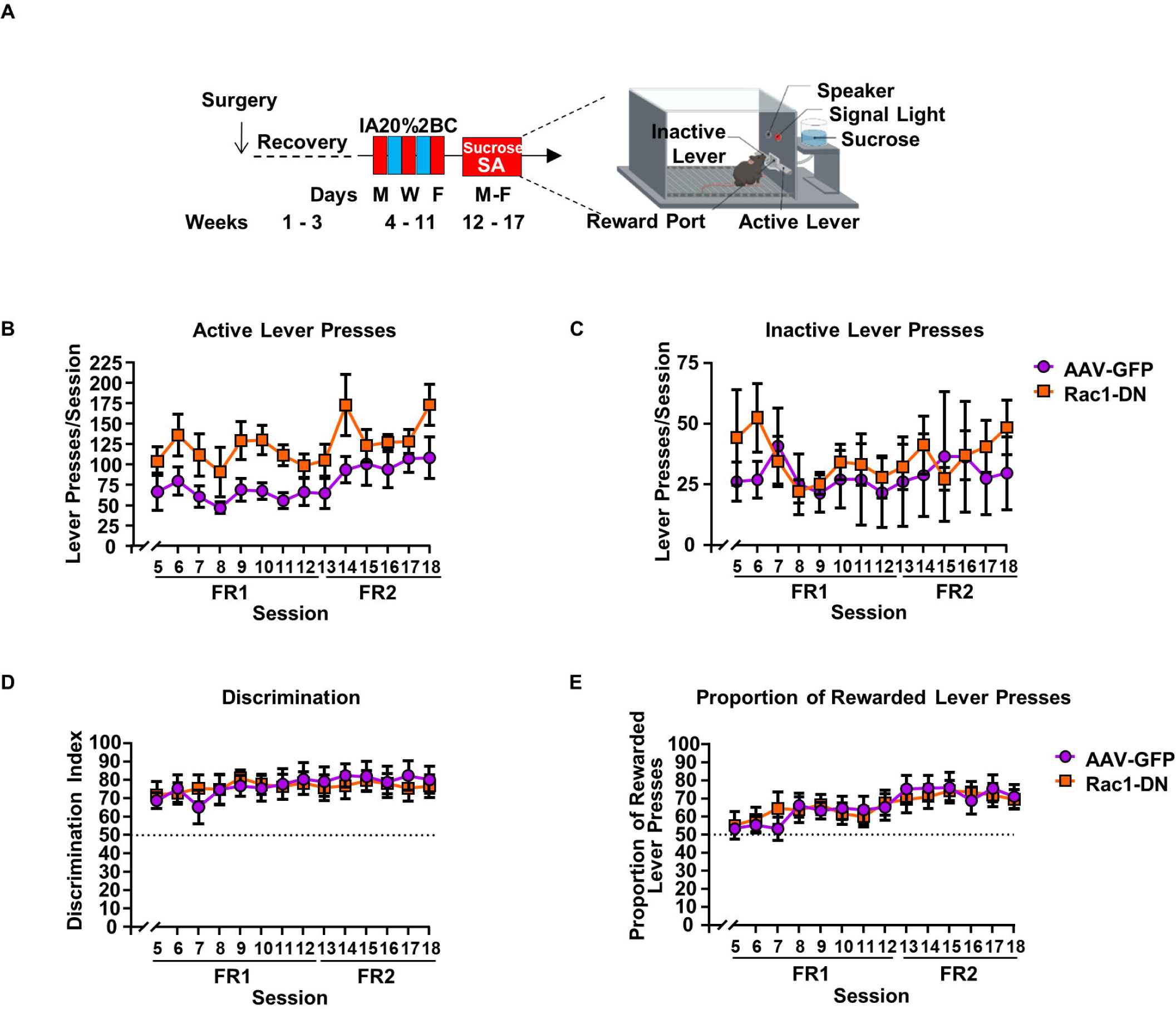
Rac1 in the DMS is not required for sucrose goal-directed learning. **(A)** Operant self-administration of 1% sucrose paradigm. Mice underwent surgery to overexpress AAV-Rac1-DN or AAV-GFP in the DMS. After 3 weeks of recovery, mice underwent IA20%2BC for 7 weeks. 1% sucrose OSA began with 1 week of training and 4 weeks of experimental tracking. Variables tracked during the session include **(B)** active lever presses and **(C)** inactive lever presses. **(D)** Discrimination and **(E)** proportion of rewarded lever presses were calculated. Data are represented as mean ± SEM and analyzed using RM Two-Way ANOVA. n = 7, 6.

## Discussion

We show herein that Rac1 signaling is activated in the DMS in response to long term excessive alcohol drinking of male mice. We further show that the consequences of alcohol-mediated Rac1/LIMK/cofilin signaling in the DMS are the formation of F-actin and the alteration of dendritic spine morphology. Finally, we present data to suggest that Rac1 in the DMS plays a role in alcohol-associated goal-directed learning. Together, our results suggest that Rac1 in the DMS plays an important role in molecular and morphological adaptations that promote alcohol-dependent learning behavior.

### Rac1 is activated in the DMS after alcohol consumption

We found that Rac1 signaling is activated specifically in the DMS of male mice in response to long-term drinking of alcohol which was observed during the 4-hour binge drinking session and was still detected after 24 hours of withdrawal. The mechanism by which alcohol activates Rac1 in the DMS is unknown, and the sequence of events is unclear, e.g. is the binge or withdrawal session that initiates the activation of Rac1? Rac1 is activated by NMDA receptor stimulation in rat cortical and hippocampal neurons (Tolias et al., 2005; Xiao et al., 2013). Previously, we found that *ex vivo* and *in vivo* exposure to, and withdrawal from, alcohol induces long-term facilitation of GluN2B-containing NMDA receptor activity specifically in the DMS (Wang et al., 2007; Wang et al., 2010). Glutamatergic tone is increased in cortical regions that project to the striatum of humans and rats during acute alcohol withdrawal (Hermann et al., 2012; Hwa et al., 2017). Therefore, it is very plausible that Rac1 is first activated during withdrawal through the activation of GluN2B receptors in the DMS, and that its activity is maintained during the alcohol drinking session. The long-lasting activation of Rac1 in the DMS in response to alcohol could be due to increased level and/or activity of one or both of its specific GEFs, Tiam1 and/or Karilin-7 (Tolias et al., 2005; Xie et al., 2007) or because of a reduction of the level and/or activity of its specific GAPs (Toure et al., 1998; Garcia-Mata et al., 2006; Peru et al., 2012; Nakamura, 2013). Another plausible explanation for alcohol-mediated Rac1 activation in the DMS is that withdrawal from IA20%2BC might trigger activation of a DMS-specific circuit by activating only neurons projecting to the DMS. For example, Ma et al. observed an increase in glutamatergic transmission from mPFC neurons projecting to the DMS in alcohol-but not water-drinking rats (Ma et al., 2017). In addition, chronic IA20%2BC drinking was shown to alter instrumental learning by affecting the thalamus-to-DMS circuit (Ma et al., 2022). Further work is required to unravel the specificity of inputs and their interaction with alcohol.

Furthermore, ∼95% of the DMS MSNs express either dopamine D1 receptors (D1) or dopamine D2 receptors (D2) (Gerfen and Surmeier, 2011; Calabresi et al., 2014; Cheng et al., 2017). Alcohol increases the complexity of dendritic branching and maturation of dendritic spines selectively in D1 but not D2 DMS MSNs (Wang et al., 2015). Since the inhibition of endogenous Rac1 attenuates alcohol-dependent re-structuring of DMS MSNs, we speculate that alcohol-mediated Rac1 signaling is specifically localized to D1 neurons. This possibility will be determined in future studies.

Interestingly, we did not detect Rac1 activation in response to alcohol drinking in the DMS of female mice. While Rac1 is not activated in the DMS of female mice in response to alcohol drinking, this does not exclude Rac1 from being activated by alcohol in other brain regions, such as the amygdala or hippocampus, where Rac1 is involved in learning and memory (Martinez et al., 2007; Haditsch et al., 2009; Wu et al., 2014; Gao et al., 2015). Further investigation is required to unravel the mechanisms controlling sex-specificity in Rac1 signaling.

### Rac1 in the DMS promotes alcohol-dependent morphological adaptations

We previously showed that the complexity of dendritic branching is increased by alcohol (Wang et al., 2015; Laguesse et al., 2018). Here, we replicated the data of both studies and showed that Rac1 is responsible for DMS MSNs dendritic complexity. However, because Rac1-DN significantly decreased dendritic arborization in both water- and alcohol-drinking animals, we cannot definitively conclude that Rac1 contributes to alcohol-dependent alteration of dendritic tree complexity, but based on the finding that Rac1 inhibition significantly alters F-actin content, this possibility is highly likely.

Rac1 is involved in the maturation and morphogenesis of dendritic spines (Nakayama et al., 2000; Tashiro et al., 2000; Pennucci et al., 2019; Costa et al., 2020). In addition to the DMS, (Wang et al., 2015; Laguesse et al., 2018) and data herein, alcohol exposure affects dendritic spine morphology in cortical regions such as the medial prefrontal cortex (mPFC) (Cannady et al., 2021), and the orbitofrontal cortex (OFC) (McGuier et al., 2015). Therefore, it is plausible that the activation of Rac1 may also be the molecular mediator of alcohol-dependent structural plasticity in cortical regions.

Interestingly, Rac1 was reported to play a role in cocaine-dependent neuroadaptations in other striatal regions. Specifically, Dietz et al. reported that cocaine-dependent increase in NAc thin spine density was mediated by Rac1 (Dietz et al., 2012), whereas Li et al. showed that the cocaine-dependent increase of Rac1 signaling in the dorsal striatum leads to dendritic spine maturation (Li et al., 2015). Cocaine also increases actin cycling through LIMK/cofilin signaling in the NAc (Toda et al., 2006). Tu et al. observed that decreased Rac1 signaling is required for METH-mediated spine density and maturation in the NAc (Tu et al., 2019). Here, the increase in dendritic maturation after alcohol drinking in the DMS is shown to be Rac1-dependent. These findings suggest that drugs of abuse are altering Rac1 signaling in a drug- and subregion-dependent manner.

### Rac1 and alcohol-associated behaviors

We found that Rac1 in the DMS does not play a role in voluntary drinking behavior. It is not unusual that alcohol-mediated signaling controls a specific behavior without affecting voluntary drinking. For example, inhibition of mTORC1 activity in the OFC attenuates habitual alcohol seeking without affecting IA20%2BC consumption (Morisot et al., 2019b). Furthermore, lack of toll-like receptor 4 (TLR4) does not alter alcohol consumption or preference (Blednov et al., 2017b), but does significantly reduce the duration of loss of righting reflex and recovery from alcohol-induced motor incoordination (Blednov et al., 2017a). Although Rac1 signaling in the DMS does not mediate voluntary drinking behavior, its activation may be involved in other alcohol-related behaviors.

The DMS primarily mediates goal-directed behaviors (Yin et al., 2005; Balleine and O’Doherty, 2010; Everitt and Robbins, 2013; Gremel and Costa, 2013), which play a major role in the addiction cycle (Hogarth, 2020). Here, we showed that Rac1 inhibition in the DMS impairs active lever discrimination and leads to a reduction in the proportion of lever presses leading to a reward, suggesting a failure to learn the association between an active lever press and alcohol reward. The elevated inactive lever presses are unlikely to have resulted from increased locomotor activity, as Rac1 inhibition did not affect locomotion. However, we found that AAV-Rac1-DN mice are insensitive to contingency degradation, implying a diminished capacity to update their action when the action-outcome association is degraded. Therefore, Rac1 in the DMS is required for alcohol-specific goal-directed action-outcome contingency or outcome value learning. While the debate persists regarding the role of goal-directed vs. habitual behavior in addiction, a large body of evidence suggest that drug seeking is driven by goal-directed behavior (Hogarth, 2020; Vandaele and Ahmed, 2021). The loss of goal-directed behavior in response to Rac1 inhibition in the DMS does not necessarily imply a transition to habitual behavior, which requires additional tests such as assessment of stimulus-response association (Turner and Balleine, 2024). Further work is required to investigate Rac1’s role in goal-directed behavior.

Interestingly, this learning deficit is specific for alcohol, as Rac1 inhibition in the DMS did not affect the learning of active lever association with sucrose reward. This phenotype could be the result of disrupting the Rac1 signaling reinforcing active lever preference (“GO”) or the avoidance of undesired action (“NO-GO”) (Cheng et al., 2017). The normal physiological role of Rac1 in the DMS is unknown. Rac1 has been linked with learning and memory in other regions such as the hippocampus, where it is required for spatial learning (Haditsch et al., 2009). Rac1 in the hippocampus also mediates reversible forgetting (Lv et al., 2019), contributing to the long-term maintenance of memory behaviors like contextual-fear conditioning and object recognition (Lv et al., 2019).

To our knowledge, this is the first study unveiling a molecular mechanism involved in alcohol-specific goal-directed learning. Prior studies have investigated the mechanisms of goal-directed learning in the context of food and sucrose seeking in the rodent striatum (Hart et al., 2018; Matamales et al., 2020; Peak et al., 2020). For example, using chemogenetics, Peak et al. found that D1R MSNs in the posterior DMS were critical for sucrose goal-directed learning (Peak et al., 2020), and Matamales et al. have shown that striatal-dependent goal-directed learning for food involves ensembles of D1R MSNs, controlled and modified by D2R MSNs (Matamales et al., 2020). It is tempting to speculate that specific sucrose or alcohol ensembles exist within the DMS. With Rac1 signaling contributing to alcohol but not sucrose self-administration, our results provide support to the notion that natural reward and alcohol drug reward mechanisms are different (Alhadeff et al., 2019).

In summary, in this study we unraveled that alcohol drinking increases Rac1 signaling in the DMS, leading to the alcohol-mediated maturation of dendritic spines and to alcohol-dependent goal-directed behavior learning processes. As goal-directed behavior is a major contributor to the cycle of addiction (Hogarth, 2020), unraveling the molecular foundation of goal-directed behavior, as explored in this study, bears profound importance for both basic science and the discovery of novel medications aimed at treating AUD.

## Author Contributions

DR conceived the project. ZWH, YE, and DR designed the experiments. ZWH, KP, AS, SL and YE conducted the experiments and analyzed the data. CS provided technical support. ZWH, YE, and DR wrote the manuscript.

## Acknowledgements

**This study was funded by** National Institute of Alcohol Abuse and Alcoholism R01 AA027682 (DR). BioRender.com was used to create some of the figures. ZWH was partially funded by NIH GM007175.

## References

Alhadeff AL, Goldstein N, Park O, Klima ML, Vargas A, Betley JN (2019) Natural and Drug Rewards Engage Distinct Pathways that Converge on Coordinated Hypothalamic and Reward Circuits. Neuron 103:891–908 e896.

Andrianantoandro E, Pollard TD (2006) Mechanism of actin filament turnover by severing and nucleation at different concentrations of ADF/cofilin. Mol Cell 24:13–23.

Balleine BW, O’Doherty JP (2010) Human and rodent homologies in action control: corticostriatal determinants of goal-directed and habitual action. Neuropsychopharmacology 35:48–69.

Bamburg JR (1999) Proteins of the ADF/cofilin family: essential regulators of actin dynamics. Annu Rev Cell Dev Biol 15:185–230.

Basu S, Lamprecht R (2018) The Role of Actin Cytoskeleton in Dendritic Spines in the Maintenance of Long-Term Memory. Front Mol Neurosci 11:143.

Bernard O (2007) Lim kinases, regulators of actin dynamics. Int J Biochem Cell Biol 39:1071–1076.

Blednov YA, Black M, Benavidez JM, Da Costa A, Mayfield J, Harris RA (2017a) Sedative and Motor Incoordination Effects of Ethanol in Mice Lacking CD14, TLR2, TLR4, or MyD88. Alcohol Clin Exp Res 41:531–540.

Blednov YA, Black M, Chernis J, Da Costa A, Mayfield J, Harris RA (2017b) Ethanol Consumption in Mice Lacking CD14, TLR2, TLR4, or MyD88. Alcohol Clin Exp Res 41:516–530.

Bokoch GM (2003) Biology of the p21-activated kinases. Annu Rev Biochem 72:743–781.

Bosco EE, Mulloy JC, Zheng Y (2009) Rac1 GTPase: a “Rac” of all trades. Cell Mol Life Sci 66:370–374.

Calabresi P, Picconi B, Tozzi A, Ghiglieri V, Di Filippo M (2014) Direct and indirect pathways of basal ganglia: a critical reappraisal. Nat Neurosci 17:1022–1030.

Cannady R, Nguyen T, Padula AE, Rinker JA, Lopez MF, Becker HC, Woodward JJ, Mulholland PJ (2021) Interaction of chronic intermittent ethanol and repeated stress on structural and functional plasticity in the mouse medial prefrontal cortex. Neuropharmacology 182:108396.

Chazeau A, Giannone G (2016) Organization and dynamics of the actin cytoskeleton during dendritic spine morphological remodeling. Cell Mol Life Sci 73:3053–3073.

Cheng Y, Huang CCY, Ma T, Wei X, Wang X, Lu J, Wang J (2017) Distinct Synaptic Strengthening of the Striatal Direct and Indirect Pathways Drives Alcohol Consumption. Biol Psychiatry 81:918–929.

Chin SM, Jansen S, Goode BL (2016) TIRF microscopy analysis of human Cof1, Cof2, and ADF effects on actin filament severing and turnover. J Mol Biol 428:1604–1616.

Corbetta S, Gualdoni S, Ciceri G, Monari M, Zuccaro E, Tybulewicz VL, de Curtis I (2009) Essential role of Rac1 and Rac3 GTPases in neuronal development. FASEB J 23:1347–1357.

Costa JF, Dines M, Lamprecht R (2020) The Role of Rac GTPase in Dendritic Spine Morphogenesis and Memory. Front Synaptic Neurosci 12:12.

Dietz DM et al. (2012) Rac1 is essential in cocaine-induced structural plasticity of nucleus accumbens neurons. Nat Neurosci 15:891–896.

Dolan RJ, Dayan P (2013) Goals and habits in the brain. Neuron 80:312–325.

Edwards DC, Sanders LC, Bokoch GM, Gill GN (1999) Activation of LIM-kinase by Pak1 couples Rac/Cdc42 GTPase signalling to actin cytoskeletal dynamics. Nat Cell Biol 1:253–259.

Everitt BJ, Robbins TW (2013) From the ventral to the dorsal striatum: devolving views of their roles in drug addiction. Neurosci Biobehav Rev 37:1946–1954.

Francis TC, Gaynor A, Chandra R, Fox ME, Lobo MK (2019) The Selective RhoA Inhibitor Rhosin Promotes Stress Resiliency Through Enhancing D1-Medium Spiny Neuron Plasticity and Reducing Hyperexcitability. Biol Psychiatry 85:1001–1010.

Gao Q, Yao W, Wang J, Yang T, Liu C, Tao Y, Chen Y, Liu X, Ma L (2015) Post-training activation of Rac1 in the basolateral amygdala is required for the formation of both short-term and long-term auditory fear memory. Front Mol Neurosci 8:65.

Garcia-Mata R, Wennerberg K, Arthur WT, Noren NK, Ellerbroek SM, Burridge K (2006) Analysis of activated GAPs and GEFs in cell lysates. Methods Enzymol 406:425–437.

Gerfen CR, Surmeier DJ (2011) Modulation of striatal projection systems by dopamine. Annu Rev Neurosci 34:441–466.

Golden SA, Christoffel DJ, Heshmati M, Hodes GE, Magida J, Davis K, Cahill ME, Dias C, Ribeiro E, Ables JL, Kennedy PJ, Robison AJ, Gonzalez-Maeso J, Neve RL, Turecki G, Ghose S, Tamminga CA, Russo SJ (2013) Epigenetic regulation of RAC1 induces synaptic remodeling in stress disorders and depression. Nat Med 19:337–344.

Gremel CM, Costa RM (2013) Orbitofrontal and striatal circuits dynamically encode the shift between goal-directed and habitual actions. Nat Commun 4:2264.

Haditsch U, Anderson MP, Freewoman J, Cord B, Babu H, Brakebusch C, Palmer TD (2013) Neuronal Rac1 is required for learning-evoked neurogenesis. J Neurosci 33:12229–12241.

Haditsch U, Leone DP, Farinelli M, Chrostek-Grashoff A, Brakebusch C, Mansuy IM, McConnell SK, Palmer TD (2009) A central role for the small GTPase Rac1 in hippocampal plasticity and spatial learning and memory. Mol Cell Neurosci 41:409–419.

Hart G, Bradfield LA, Balleine BW (2018) Prefrontal Corticostriatal Disconnection Blocks the Acquisition of Goal-Directed Action. J Neurosci 38:1311–1322.

Hermann D, Weber-Fahr W, Sartorius A, Hoerst M, Frischknecht U, Tunc-Skarka N, Perreau-Lenz S, Hansson AC, Krumm B, Kiefer F, Spanagel R, Mann K, Ende G, Sommer WH (2012) Translational magnetic resonance spectroscopy reveals excessive central glutamate levels during alcohol withdrawal in humans and rats. Biol Psychiatry 71:1015–1021.

Hogarth L (2020) Addiction is driven by excessive goal-directed drug choice under negative affect: translational critique of habit and compulsion theory. Neuropsychopharmacology 45:720–735.

Honkura N, Matsuzaki M, Noguchi J, Ellis-Davies GC, Kasai H (2008) The subspine organization of actin fibers regulates the structure and plasticity of dendritic spines. Neuron 57:719–729.

Hotulainen P, Hoogenraad CC (2010) Actin in dendritic spines: connecting dynamics to function. J Cell Biol 189:619–629.

Hwa L, Besheer J, Kash T (2017) Glutamate plasticity woven through the progression to alcohol use disorder: a multi-circuit perspective. F1000Res 6:298.

Kanellos G, Frame MC (2016) Cellular functions of the ADF/cofilin family at a glance. Journal of Cell Science 129:3211–3218.

Kasai H, Matsuzaki M, Noguchi J, Yasumatsu N, Nakahara H (2003) Structure-stability-function relationships of dendritic spines. Trends Neurosci 26:360–368.

Koob G, Kreek MJ (2007) Stress, dysregulation of drug reward pathways, and the transition to drug dependence. Am J Psychiatry 164:1149–1159.

Kravitz AV, Kreitzer AC (2012) Striatal mechanisms underlying movement, reinforcement, and punishment. Physiology (Bethesda) 27:167–177.

Laguesse S, Morisot N, Phamluong K, Sakhai SA, Ron D (2018) mTORC2 in the dorsomedial striatum of mice contributes to alcohol-dependent F-Actin polymerization, structural modifications, and consumption. Neuropsychopharmacology 43:1539–1547.

Laguesse S, Morisot N, Shin JH, Liu F, Adrover MF, Sakhai SA, Lopez MF, Phamluong K, Griffin WC, 3rd, Becker HC, Bender KJ, Alvarez VA, Ron D (2017) Prosapip1-Dependent Synaptic Adaptations in the Nucleus Accumbens Drive Alcohol Intake, Seeking, and Reward. Neuron 96:145–159 e148.

Li J, Zhang L, Chen Z, Xie M, Huang L, Xue J, Liu Y, Liu N, Guo F, Zheng Y, Kong J, Zhang L, Zhang L (2015) Cocaine activates Rac1 to control structural and behavioral plasticity in caudate putamen. Neurobiol Dis 75:159–176.

Lv L, Liu Y, Xie J, Wu Y, Zhao J, Li Q, Zhong Y (2019) Interplay between alpha2-chimaerin and Rac1 activity determines dynamic maintenance of long-term memory. Nat Commun 10:5313.

Ma T, Barbee B, Wang X, Wang J (2017) Alcohol induces input-specific aberrant synaptic plasticity in the rat dorsomedial striatum. Neuropharmacology 123:46–54.

Ma T, Huang Z, Xie X, Cheng Y, Zhuang X, Childs MJ, Gangal H, Wang X, Smith LN, Smith RJ, Zhou Y, Wang J (2022) Chronic alcohol drinking persistently suppresses thalamostriatal excitation of cholinergic neurons to impair cognitive flexibility. J Clin Invest 132.

Maciver SK, Hussey PJ (2002) The ADF/cofilin family: actin-remodeling proteins. Genome Biol 3:reviews3007.

Martinez LA, Klann E, Tejada-Simon MV (2007) Translocation and activation of Rac in the hippocampus during associative contextual fear learning. Neurobiol Learn Mem 88:104–113.

Martins JS, Joyner KJ, McCarthy DM, Morris DH, Patrick CJ, Bartholow BD (2022) Differential brain responses to alcohol-related and natural rewards are associated with alcohol use and problems: Evidence for reward dysregulation. Addict Biol 27:e13118.

Matamales M, McGovern AE, Mi JD, Mazzone SB, Balleine BW, Bertran-Gonzalez J (2020) Local D2- to D1-neuron transmodulation updates goal-directed learning in the striatum. Science 367:549–555.

McGuier NS, Padula AE, Lopez MF, Woodward JJ, Mulholland PJ (2015) Withdrawal from chronic intermittent alcohol exposure increases dendritic spine density in the lateral orbitofrontal cortex of mice. Alcohol 49:21–27.

Mira JP, Benard V, Groffen J, Sanders LC, Knaus UG (2000) Endogenous, hyperactive Rac3 controls proliferation of breast cancer cells by a p21-activated kinase-dependent pathway. Proc Natl Acad Sci U S A 97:185–189.

Morisot N, Berger AL, Phamluong K, Cross A, Ron D (2019a) The Fyn kinase inhibitor, AZD0530, suppresses mouse alcohol self-administration and seeking. Addict Biol 24:1227–1234.

Morisot N, Phamluong K, Ehinger Y, Berger AL, Moffat JJ, Ron D (2019b) mTORC1 in the orbitofrontal cortex promotes habitual alcohol seeking. Elife 8.

Nakamura F (2013) FilGAP and its close relatives: a mediator of Rho-Rac antagonism that regulates cell morphology and migration. Biochem J 453:17–25.

Nakayama AY, Harms MB, Luo L (2000) Small GTPases Rac and Rho in the maintenance of dendritic spines and branches in hippocampal pyramidal neurons. J Neurosci 20:5329–5338.

Nall RW, Heinsbroek JA, Nentwig TB, Kalivas PW, Bobadilla AC (2021) Circuit selectivity in drug versus natural reward seeking behaviors. J Neurochem 157:1450–1472.

Nguyen LK, Kholodenko BN, von Kriegsheim A (2018) Rac1 and RhoA: Networks, loops and bistability. Small GTPases 9:316–321.

Pavlov D, Muhlrad A, Cooper J, Wear M, Reisler E (2007) Actin filament severing by cofilin. J Mol Biol 365:1350–1358.

Peak J, Chieng B, Hart G, Balleine BW (2020) Striatal direct and indirect pathway neurons differentially control the encoding and updating of goal-directed learning. Elife 9.

Pennucci R, Gucciardi I, de Curtis I (2019) Rac1 and Rac3 GTPases differently influence the morphological maturation of dendritic spines in hippocampal neurons. PLoS One 14:e0220496.

Peru YCdPRL, Acevedo SF, Rodan AR, Chang LY, Eaton BA, Rothenfluh A (2012) Adult neuronal Arf6 controls ethanol-induced behavior with Arfaptin downstream of Rac1 and RhoGAP18B. J Neurosci 32:17706–17713.

Ratner N, Mahler HR (1983) Structural organization of filamentous proteins in postsynaptic density. Biochemistry 22:2446–2453.

Reijnders MRF et al. (2017) RAC1 Missense Mutations in Developmental Disorders with Diverse Phenotypes. Am J Hum Genet 101:466–477.

Ridley AJ (2006) Rho GTPases and actin dynamics in membrane protrusions and vesicle trafficking. Trends Cell Biol 16:522–529.

Ridley AJ, Paterson HF, Johnston CL, Diekmann D, Hall A (1992) The small GTP-binding protein rac regulates growth factor-induced membrane ruffling. Cell 70:401–410.

Schubert V, Dotti CG (2007) Transmitting on actin: synaptic control of dendritic architecture. J Cell Sci 120:205–212.

Scott RW, Olson MF (2007) LIM kinases: function, regulation and association with human disease. J Mol Med (Berl) 85:555–568.

Shan Q, Ge M, Christie MJ, Balleine BW (2014) The acquisition of goal-directed actions generates opposing plasticity in direct and indirect pathways in dorsomedial striatum. J Neurosci 34:9196–9201.

Sholl DA (1953) Dendritic organization in the neurons of the visual and motor cortices of the cat. J Anat 87:387–406.

Singer BF, Fadanelli M, Kawa AB, Robinson TE (2018) Are Cocaine-Seeking “Habits” Necessary for the Development of Addiction-Like Behavior in Rats? J Neurosci 38:60–73.

Swanson AM, DePoy LM, Gourley SL (2017) Inhibiting Rho kinase promotes goal-directed decision making and blocks habitual responding for cocaine. Nat Commun 8:1861.

Tashiro A, Minden A, Yuste R (2000) Regulation of dendritic spine morphology by the rho family of small GTPases: antagonistic roles of Rac and Rho. Cereb Cortex 10:927–938.

Tejada-Simon MV (2015) Modulation of actin dynamics by Rac1 to target cognitive function. J Neurochem 133:767–779.

Toda S, Shen HW, Peters J, Cagle S, Kalivas PW (2006) Cocaine increases actin cycling: effects in the reinstatement model of drug seeking. J Neurosci 26:1579–1587.

Tolias KF, Bikoff JB, Burette A, Paradis S, Harrar D, Tavazoie S, Weinberg RJ, Greenberg ME (2005) The Rac1-GEF Tiam1 couples the NMDA receptor to the activity-dependent development of dendritic arbors and spines. Neuron 45:525–538.

Toure A, Dorseuil O, Morin L, Timmons P, Jegou B, Reibel L, Gacon G (1998) MgcRacGAP, a new human GTPase-activating protein for Rac and Cdc42 similar to Drosophila rotundRacGAP gene product, is expressed in male germ cells. J Biol Chem 273:6019–6023.

Tu G, Ying L, Ye L, Zhao J, Liu N, Li J, Liu Y, Zhu M, Wu Y, Xiao B, Guo H, Guo F, Wang H, Zhang L, Zhang L (2019) Dopamine D(1) and D(2) Receptors Differentially Regulate Rac1 and Cdc42 Signaling in the Nucleus Accumbens to Modulate Behavioral and Structural Plasticity After Repeated Methamphetamine Treatment. Biol Psychiatry 86:820–835.

Turner KM, Balleine BW (2024) Stimulus control of habits: Evidence for both stimulus specificity and devaluation insensitivity in a dual-response task. J Exp Anal Behav 121:52–61.

Van Aelst L, D’Souza-Schorey C (1997) Rho GTPases and signaling networks. Genes Dev 11:2295–2322.

Vandaele Y, Ahmed SH (2021) Habit, choice, and addiction. Neuropsychopharmacology 46:689–698.

Vetter IR, Wittinghofer A (2001) The guanine nucleotide-binding switch in three dimensions. Science 294:1299–1304.

Wang J, Lanfranco MF, Gibb SL, Yowell QV, Carnicella S, Ron D (2010) Long-lasting adaptations of the NR2B-containing NMDA receptors in the dorsomedial striatum play a crucial role in alcohol consumption and relapse. J Neurosci 30:10187–10198.

Wang J, Cheng Y, Wang X, Roltsch Hellard E, Ma T, Gil H, Ben Hamida S, Ron D (2015) Alcohol Elicits Functional and Structural Plasticity Selectively in Dopamine D1 Receptor-Expressing Neurons of the Dorsomedial Striatum. J Neurosci 35:11634–11643.

Wang J, Carnicella S, Phamluong K, Jeanblanc J, Ronesi JA, Chaudhri N, Janak PH, Lovinger DM, Ron D (2007) Ethanol induces long-term facilitation of NR2B-NMDA receptor activity in the dorsal striatum: implications for alcohol drinking behavior. J Neurosci 27:3593–3602.

Wang W, Ju YY, Zhou QX, Tang JX, Li M, Zhang L, Kang S, Chen ZG, Wang YJ, Ji H, Ding YQ, Xu L, Liu JG (2017) The Small GTPase Rac1 Contributes to Extinction of Aversive Memories of Drug Withdrawal by Facilitating GABA(A) Receptor Endocytosis in the vmPFC. J Neurosci 37:7096–7110.

Wang X, Liu D, Wei F, Li Y, Wang X, Li L, Wang G, Zhang S, Zhang L (2020) Stress-Sensitive Protein Rac1 and Its Involvement in Neurodevelopmental Disorders. Neural Plast 2020:8894372.

Warnault V, Darcq E, Morisot N, Phamluong K, Wilbrecht L, Massa SM, Longo FM, Ron D (2016) The BDNF Valine 68 to Methionine Polymorphism Increases Compulsive Alcohol Drinking in Mice That Is Reversed by Tropomyosin Receptor Kinase B Activation. Biol Psychiatry 79:463–473.

Wong KW, Mohammadi S, Isberg RR (2006) Disruption of RhoGDI and RhoA regulation by a Rac1 specificity switch mutant. J Biol Chem 281:40379–40388.

Worthylake DK, Rossman KL, Sondek J (2000) Crystal structure of Rac1 in complex with the guanine nucleotide exchange region of Tiam1. Nature 408:682–688.

Wu P, Ding ZB, Meng SQ, Shen HW, Sun SC, Luo YX, Liu JF, Lu L, Zhu WL, Shi J (2014) Differential role of Rac in the basolateral amygdala and cornu ammonis 1 in the reconsolidation of auditory and contextual Pavlovian fear memory in rats. Psychopharmacology (Berl) 231:2909–2919.

Xiao L, Hu C, Yang W, Guo D, Li C, Shen W, Liu X, Aijun H, Dan W, He C (2013) NMDA receptor couples Rac1-GEF Tiam1 to direct oligodendrocyte precursor cell migration. Glia 61:2078–2099.

Xie Z, Srivastava DP, Photowala H, Kai L, Cahill ME, Woolfrey KM, Shum CY, Surmeier DJ, Penzes P (2007) Kalirin-7 controls activity-dependent structural and functional plasticity of dendritic spines. Neuron 56:640–656.

Yang N, Higuchi O, Ohashi K, Nagata K, Wada A, Kangawa K, Nishida E, Mizuno K (1998) Cofilin phosphorylation by LIM-kinase 1 and its role in Rac-mediated actin reorganization. Nature 393:809–812.

Yin HH, Ostlund SB, Knowlton BJ, Balleine BW (2005) The role of the dorsomedial striatum in instrumental conditioning. Eur J Neurosci 22:513–523.

Zhang C, Liu J, Zhao Y, Yue X, Zhu Y, Wang X, Wu H, Blanco F, Li S, Bhanot G, Haffty BG, Hu W, Feng Z (2016) Glutaminase 2 is a novel negative regulator of small GTPase Rac1 and mediates p53 function in suppressing metastasis. Elife 5:e10727.

Zhang H, Ben Zablah Y, Zhang H, Jia Z (2021) Rho Signaling in Synaptic Plasticity, Memory, and Brain Disorders. Front Cell Dev Biol 9:729076.

